# Estrogen Regulates the Satellite Cell Compartment in Females

**DOI:** 10.1101/331777

**Authors:** Brittany C. Collins, Robert W. Arpke, Alexie A. Larson, Cory W. Baumann, Christine A Cabelka, Nardina L. Nash, Hanna-Kaarina Juppi, Eija K. Laakkonen, Sarianna Sipilä, Vuokko Kovanen, Espen E. Spangenburg, Michael Kyba, Dawn A. Lowe

**Affiliations:** Divisions of Rehabilitation Science and Physical Therapy, Department of Rehabilitation Medicine, Medical School, University of Minnesota, Minneapolis, Minnesota, USA; Lillehei Heart Institute and Department of Pediatrics, Medical School, University of Minnesota, Minneapolis, Minnesota, USA; Department of Integrative Biology and Physiology, Medical School, University of Minnesota, Minneapolis, Minnesota, USA; Gerontology Research Center, Faculty of Sport and Health Sciences, University of Jyväskylä, Jyväskylä, Finland; East Carolina Diabetes and Obesity Institute, Department of Physiology, Brody School of Medicine, East Carolina University, Greenville, NC, USA

**Keywords:** estradiol, muscle stem cells, ovarian hormones, quiescence, skeletal muscle

## Abstract

Skeletal muscle mass, strength, and regenerative capacity decline with age, with many measures showing greater deterioration in females about the time estrogen levels decrease at menopause. Here we show that maintenance of muscle stem cells, satellite cells, as well as self-renewal and differentiation into muscle fibers, are severely compromised by estrogen deficiency. Mechanistically, by hormone replacement, use of a selective estrogen-receptor modulator (bazedoxifene), and conditional estrogen receptor knockout, we implicate 17β-estradiol and satellite cell expression of estrogen receptor *α* (ERα) and show that estrogen signaling through this receptor is necessary to prevent apoptosis of satellite cells. Early data from a biopsy study of women who transitioned from peri-to post-menopause are consistent with the loss of satellite cells coincident with the decline in estradiol in humans. Together, these results demonstrate an important role for estrogen in satellite cell maintenance and muscle regeneration in females.

## INTRODUCTION

Over the course of an individual’s life, skeletal muscle undergoes numerous injurious insults that require repair in order for function to be maintained. The maintenance and injury repair of skeletal muscle is dependent on its resident stem cell, i.e., the satellite cell, with genetic ablation of satellite cells completely abolishing the ability of skeletal muscle to regenerate following injury (Fry et al., 2015; Murphy et al., 2011; Sambasivan et al., 2011). Satellite cells are located between the sarcolemma and the basal lamina of skeletal muscle fibers where they remain in a quiescent state (Conboy and Rando, 2002; Fukada et al., 2007; Keefe et al., 2015; Kuang et al., 2007), becoming activated through external stimuli, such as a muscle injury, initiating the transition from quiescence into the myogenic program to repair damaged muscle (Conboy and Rando, 2002; Dumont et al., 2015; Hindi and Kumar, 2016; Kuang et al., 2007; Troy et al., 2012). With proliferation, satellite cells undergo asymmetric division through which a subpopulation of the daughter satellite cells do not differentiate but instead return to quiescence repopulating the satellite cell pool (i.e., self-renewal) (Kuang et al., 2007; Troy et al., 2012). The balance of this asymmetric division process is critical and necessary to ensure life-long preservation of satellite cells in skeletal muscle.

Aging diminishes the satellite cell pool (Keefe et al., 2015; Sajko et al., 2004; Verdijk et al., 2014) and as a result the regenerative capacity of skeletal muscle in aged males is impaired, compared to that of younger males (Brack et al., 2005; Carlson and Conboy, 2007; Chakkalakal et al., 2012; Keefe et al., 2015; Shefer et al., 2006), but such age-induced impairments in females is less studied. Similarly, age-associated changes in the satellite cell environment, in combination with cell intrinsic alterations, disrupt quiescence and the balance of asymmetric division, ultimately impacting satellite cell maintenance and muscle regenerative potential (Bernet et al., 2014; Conboy et al., 2005; Cosgrove et al., 2014; Sousa-Victor et al., 2014). Such results support the concept that circulatory factors, including hormones that differ between the young and old systemic environment and the activity of their subsequent signaling pathways, contribute to age-associated decrements in satellite cell maintenance and overall muscle regenerative capacity.

A well-known hormone that changes with age is estradiol, the main circulating sex hormone in adult females. Estradiol is a major regulator of not only gonadal organ development and function, it has recently been recognized for its protective effects in certain tissues (e.g., against cardiovascular disease and osteoporosis) in women prior to the menopausal transition (Deschamps et al., 2010). Serum estradiol concentration declines dramatically at the average age of 51 in women, corresponding to the time of menopause (Baber et al., 2016). Estradiol deficiency reduces skeletal muscle mass and force generation in women (Greising et al., 2009; Phillips et al., 1993; Phillips et al., 1996; Qaisar et al., 2013; Taaffe et al., 2005) and female rodents (Greising et al., 2011; Le et al., 2017; Moran et al., 2007) and prevents full recovery of strength following contraction-induced injury (Kosir et al., 2015; Rader and Faulkner, 2006). Mechanistically, it has been hypothesized that the regulation of muscle inflammation (Le et al., 2015; McClung et al., 2007; Tiidus et al., 2001) and protein synthesis (Kamanga-Sollo et al., 2010; Kosir et al., 2015; McClung et al., 2007; McClung et al., 2006) is perturbed by estradiol deficiency and may contribute to impaired recovery (Enns et al., 2008). Previous work has also indicated that exercise-induced activation of satellite cells is less effective in the absence of estradiol (Enns and Tiidus, 2008), and that androgens contribute to the regulation of juvenile satellite cells during growth (Kim et al., 2016), but it has yet to be determined if there are cell autonomous mechanisms which involve estradiol acting directly on satellite cells.

## RESULTS

### Estradiol regulates satellite cell maintenance

To directly test whether ovarian hormone deficiency affects the satellite cell compartment, we counted the total number of satellite cells in five diverse muscles from control and ovariectomized (Ovx) mice (Supplementary Fig. 1). Satellite cells (lineage negative; VCAM, alpha7 double positive) were 30-60% fewer in number in tibialis anterior (TA) muscles of Ovx than Control mice and reduction was associated with the duration of hormone deficiency (Fig. 1a). Reductions were also measured in the EDL, gastrocnemius, and diaphragm (Fig. 1a). The soleus, which is a slower and more fatigue-resistant muscle, was unaffected by hormone manipulation (Fig. 1a). We also evaluated the density of satellite cells, calculated by dividing cell number (Fig. 1a) by the wet mass of each muscle (Fig. 1b,c). Satellite cell density recapitulated the cell number declines with ovarian hormone deficiency in TA, gastrocnemius, and EDL muscles. In the diaphragm, where the decline in total number was only statistically significant at the 2 mo point, the decline in satellite cell density was statistically significant at all time points studied (Fig. 1c).

**Figure 1.**
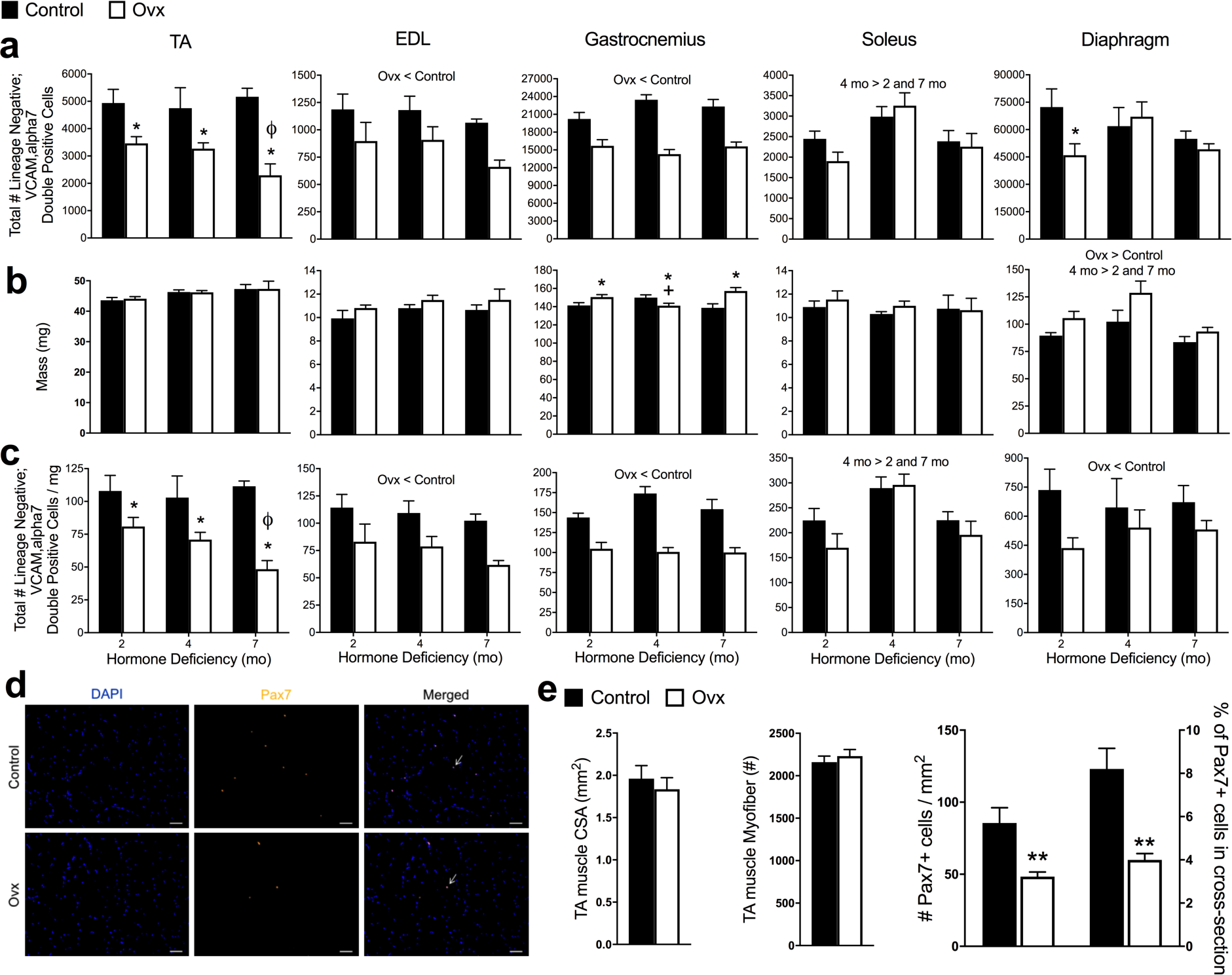
Estrogen deficiency disrupts maintenance of satellite cells in skeletal muscles of females. **(a)** Total number of satellite cells quantified by lineage negative;VCAM,alpha7 double-positive cells in five discrete muscles from Control (n=15) and ovariectomized (Ovx; n=15) mice. Muscles were harvested and analyzed 2, 4, or 7 mo after Ovx and in age-matched controls. **(b)** Muscle masses. **(c)** Total number of satellite cells normalized to muscle masses. **(d)** Satellite cells quantified by immunohistochemistry of Pax7+ cells in TA muscles from Control (n=4) and Ovx (n=4) mice at 2 mo of hormone deficiency. Arrows indicate localization of DAPI+ Pax7+double-positive cell. Scale bars = 50 µm. **(e)** TA muscle cross-sectional area and number of fibers from Control and Ovx mice (P≥0.193), and Pax7+ cells per TA muscle cross-sectional area and % Pax7+ cells in each cross-section relative to the total # of muscle fibers. Significant main effects of two-way ANOVA (P<0.05) are indicated above the bars (abc) and when interactions occurred (P<0.05), Holm–Sidak post-hoc tests are indicated by *different than Control at corresponding duration (abc), ^ϕ^ different than 2 and 4 mo Ovx (ac), and ^+^ different than 2 and 7 mo Ovx (b). **P<0.005 by student t-tests (e).

To confirm the effect of ovarian hormone deficiency on satellite cell number using an independent marker, we employed the Pax7-ZsGreen mouse model, in which quiescent satellite cells fluoresce green and differentiating cells rapidly lose fluorescence (Bosnakovski et al., 2008). ZsGreen+ cells were measured in the same five skeletal muscles of Control and Ovx Pax7-ZsGreen mice at 2 mo of hormone deficiency (Supplementary Fig. 2). Consistent with results from surface marker staining, ZsGreen+ cells were significantly lower in the TA, EDL and gastrocnemius muscles of Ovx mice, with the diaphragm trending and soleus again being less affected (Supplementary Fig. 2). To independently verify the results of FACS quantification, we also counted Pax7+ cells in immuno-stained TA muscle sections (Fig. 1d). Histological analysis showed that TA muscles from Ovx mice have ∼50% fewer satellite cells than controls (Fig. 1e).

**Figure 2.**
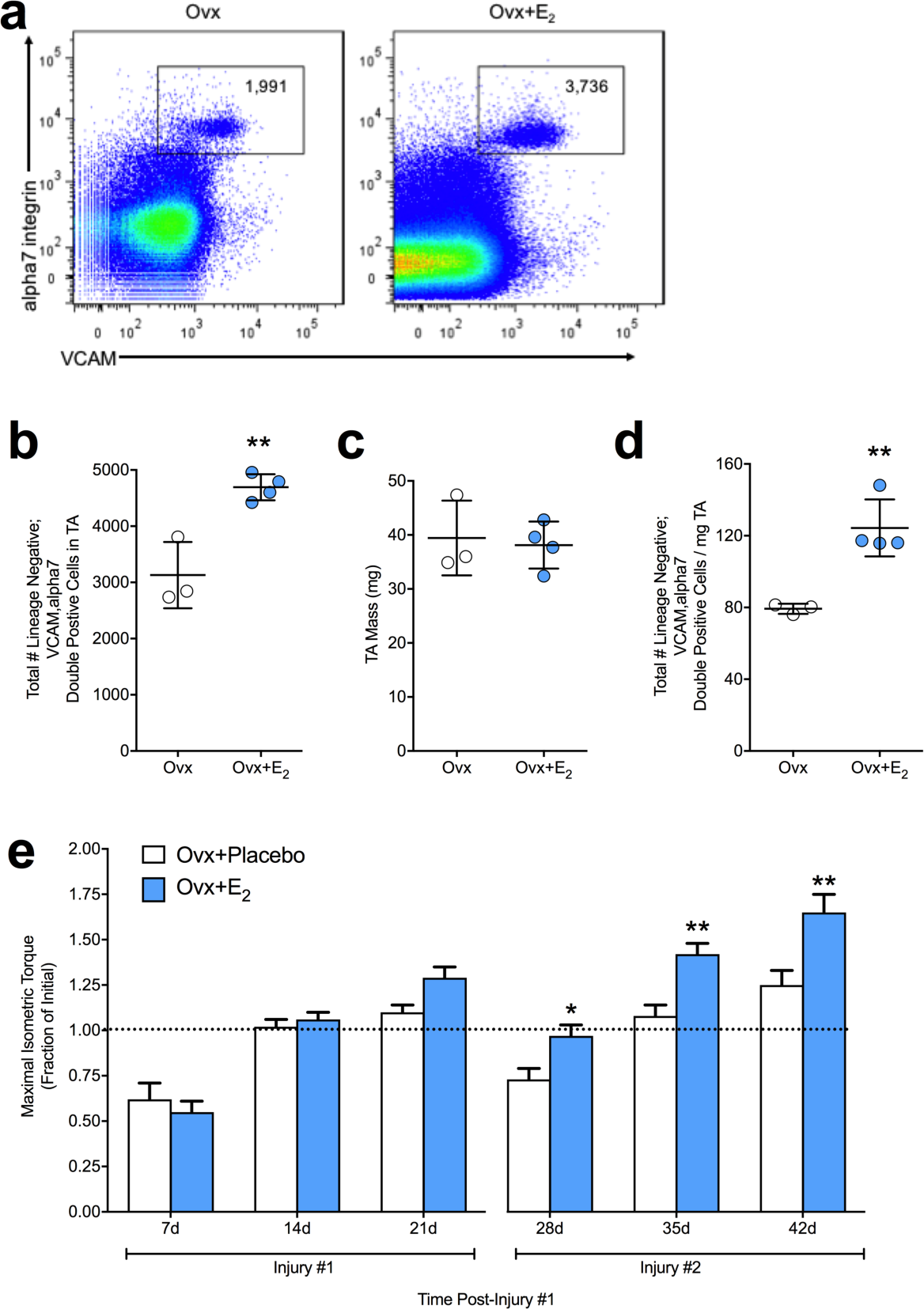
Estradiol maintains the satellite cell pool and accelerates recovery of strength. **(a)** Representative FACS plots of cells isolated from TA muscles of Ovx mice without (n=3) and with 17β-estradiol treatment (Ovx+E_2_; n=4). **(b)** Total number of satellite cells and (**cd**) number relative to TA muscle mass. (**e**) Maximal isometric torque (i.e., strength) expressed relative to pre-injury torque in Ovx mice without (Ovx+Placebo; n=6) or with 17β-estradiol treatment (Ovx+E_2_; n=8) following repeated injuries to TA muscle. * P<0.05 and **P<0.005 by student t-tests (abcd) and Holm-Sidak post hoc (e).

Ovariectomy results in systemic changes and deficiencies of multiple hormones. To determine if estradiol was the hormone responsible for affecting satellite cells, a subset of Ovx mice was treated with 17β-estradiol simultaneously with Ovx. Treatment with this specific ovarian hormone rescued satellite cell numbers, preventing depletion of the satellite cell pool (Fig. 2 a-d), thus demonstrating sufficiency for the hormone estradiol.

Given that loss of estradiol perturbs maintenance of the satellite cell pool during homeostatic conditions, we aimed to determine if this loss of satellite cell number resulted in functional consequences. We investigated if recovery of function (i.e., strength) from a chemical injury to the TA muscle was affected by the loss of estradiol (Ovx+Placebo). Following a single bout of injury, estradiol deficiency did not affect strength recovery until 21 days post-injury which resulted in a 19% lower torque strength (P=0.06; Fig. 2e). Strikingly, following a second bout of injury strength recovery was substantially lower by 23-39% at the three time-points, 28, 35, and 42 days post-injury. Treatment with estradiol (Ovx+E_2_) rescued the decrements in strength (P≤0.019; Fig. 2e). These results demonstrate that the decline in satellite cell number is associated with an impaired regenerative response.

### Estradiol deficiency impairs self-renewal and differentiation *in vivo*

The disruption of satellite cell maintenance and the impairment in strength recovery following injury that occurred with the loss of estradiol led us to next determine if self-renewal and/or differentiation were affected as these cellular processes have direct implications for replenishment of the satellite cell pool. We assayed self-renewal and differentiation *in vivo* via transplantation. Satellite cells were harvested from bulk hindlimb cell preparations from female Pax7-ZsGreen mice and 600 cells were transplanted into previously irradiated, cardiotoxin-injured TA muscles of Control or Ovx recipient syngeneic C57/BL6J mice (Fig. 3a). Satellite cell engraftment was measured by FACS using the donor ZsGreen+ marker, which is maintained only in undifferentiated Pax7+ donor-derived satellite (Fig. 3b). Estradiol deficiency of the recipient resulted in 75% lower engraftment of the satellite cell compartment compared to control recipients (Fig. 3c).

**Figure 3.**
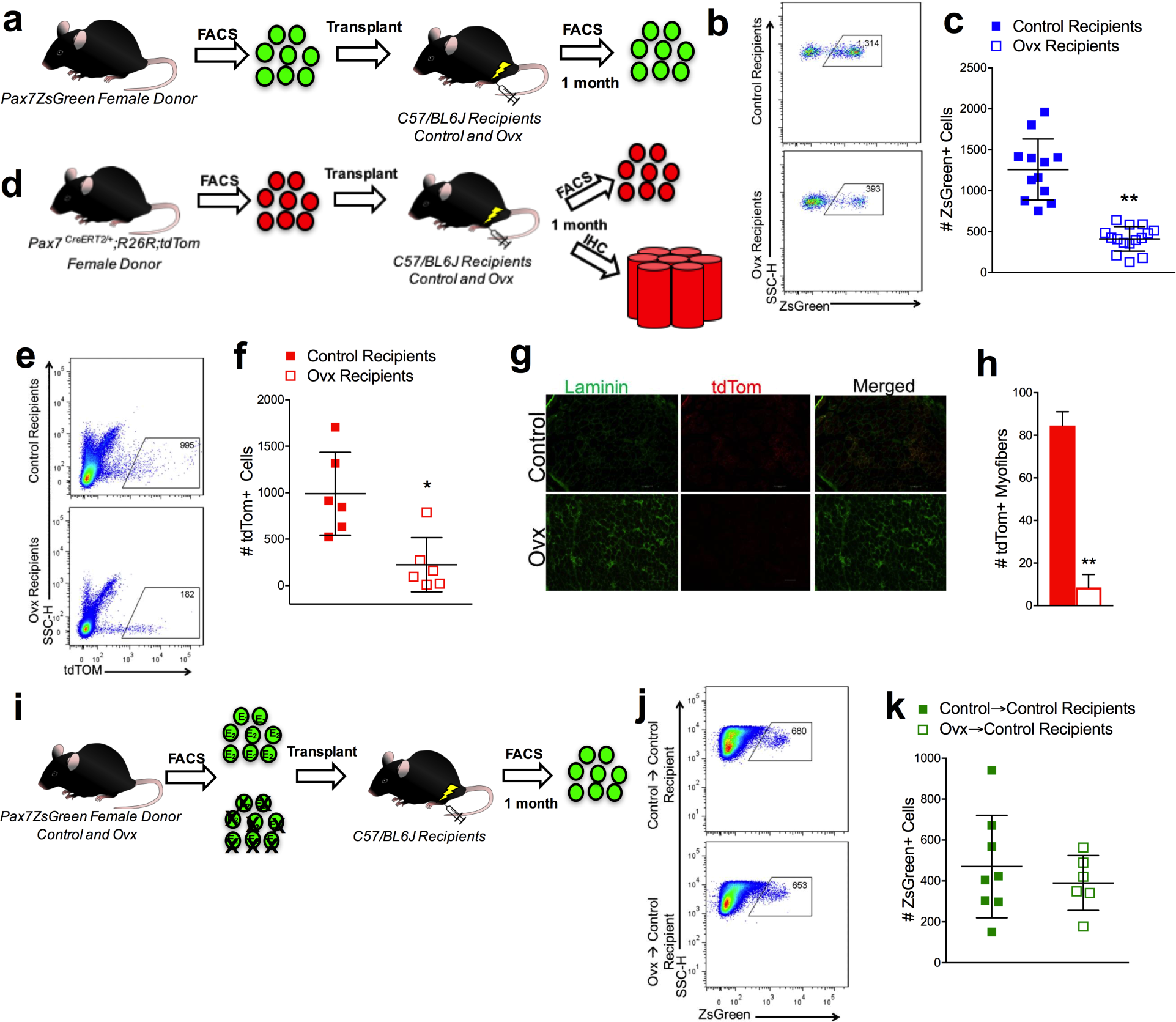
Loss of estradiol in the environment negatively affects satellite cell and fiber engraftment that can be rescued by the presence of estradiol. **(a)** Transplantation scheme for ZsGreen+ cells transplanted into Control and ovariectomized (Ovx) recipients. **(b)** Representative FACS plots of ZsGreen+ donor satellite cells in TA muscle 1 mo post-transplant in Control and Ovx recipient mice. **(c)** Total number of ZsGreen+ donor satellite cells in Control (n=12) and Ovx (n=15) recipient mice 1 mo post-transplant. **(d)** Scheme for tdTom+ cells transplanted into Control (n=6) and Ovx (n=6) recipients. **(e)** Representative FACS plots of tdTOM+ donor satellite cells in TA muscle 1 mo post-transplant in Control and Ovx recipient mice. **(f)** Total number of tdTOM+ donor satellite cells in Control (n=6) and Ovx (n=6) recipient mice 1 mo post-transplant. **(g)** Representative images of tdTom+ fibers in the engrafted region of TA muscle from Control and Ovx recipient mice. Scale bars = 100 µm. **(h)** Quantification of total number of tdTom+ fibers from Control and Ovx recipients. **(i)** Transplantation scheme for ZsGreen+ cells from Control and Ovx donors. **(j)** Representative FACS plots of ZsGreen+ Control and Ovx donor satellite cells in TA muscle 1 mo following transplant in Control recipient mice. **(k)** Total number of ZsGreen+ Control (n=8) and Ovx (n=6) donor satellite cells in Control recipient TA muscles 1 mo post-transplantation. * P<0.05 and **P<0.005 by student t-tests

To address the contribution of satellite cells to fibers, we utilized a lineage-tracing strategy (Pax7 ^CreERT2/+^;R26R^tdTom^ mice) to mark donor-derived fibers (Fig. 3d). This experiment again confirmed the deleterious effect of systemic estradiol deficiency on satellite cell engraftment (Fig. 3e,f) and further demonstrated that while satellite cells contributed robustly to fibers in muscle of control recipients, contribution to fibers was minimal when estradiol was not present (Fig. 3g,h). As a result of the reduced engraftment and differentiation potential of the satellite cells when transplanted from an estradiol-replete environment into one lacking estradiol, we next aimed to determine if satellite cells from an environment lacking estradiol (i.e., from Ovx donors) were intrinsically, irreversibly impaired, or whether they would be functional in an estrogen-replete environment. ZsGreen+ satellite cells were isolated from the TA and gastrocnemius muscles of Control and Ovx donor mice and 600 cells were transplanted into control recipients (Fig. 3i). The transfer into an estradiol-replete environment (i.e., control recipient) rendered satellite cells from Ovx mice similar to those of Controls (Fig. 3j,k).

### Apoptosis of satellite cells with estradiol deficiency

We next investigated whether the mechanism for the reduction in cell number could involve apoptosis. In non-skeletal muscle tissues, estradiol is known to protect against apoptosis (e.g., Guo et al., 2013) and immortalized myoblasts were also found to undergo less apoptosis *in vitro* in the presence of estradiol (Boland et al., 2008; La Colla et al., 2016; Vasconsuelo et al., 2010). It has been suggested that apoptosis plays an important role in skeletal muscle health, ultimately affecting strength (La Colla et al., 2015; Pallafacchina et al., 2013; Sanchez et al., 2014). However, it is important to point out that in healthy unperturbed muscle, apoptotic cells are from other lineages and that rates of apoptosis of quiescent satellite cells during homeostatic conditions are nearly zero (Fry et al., 2016; Hirai et al., 2010; Shea et al., 2010). First, we measured overall apoptosis *in vivo* by identifying TUNEL+ cells in the TA muscles of Control and Ovx mice (Fig. 3a). Apoptotic cells were found in muscle from both groups, with estradiol-deficient muscles having 3.7-fold more TUNEL+ cells compared to Controls (P=0.08; Fig. 4b).

**Figure 4.**
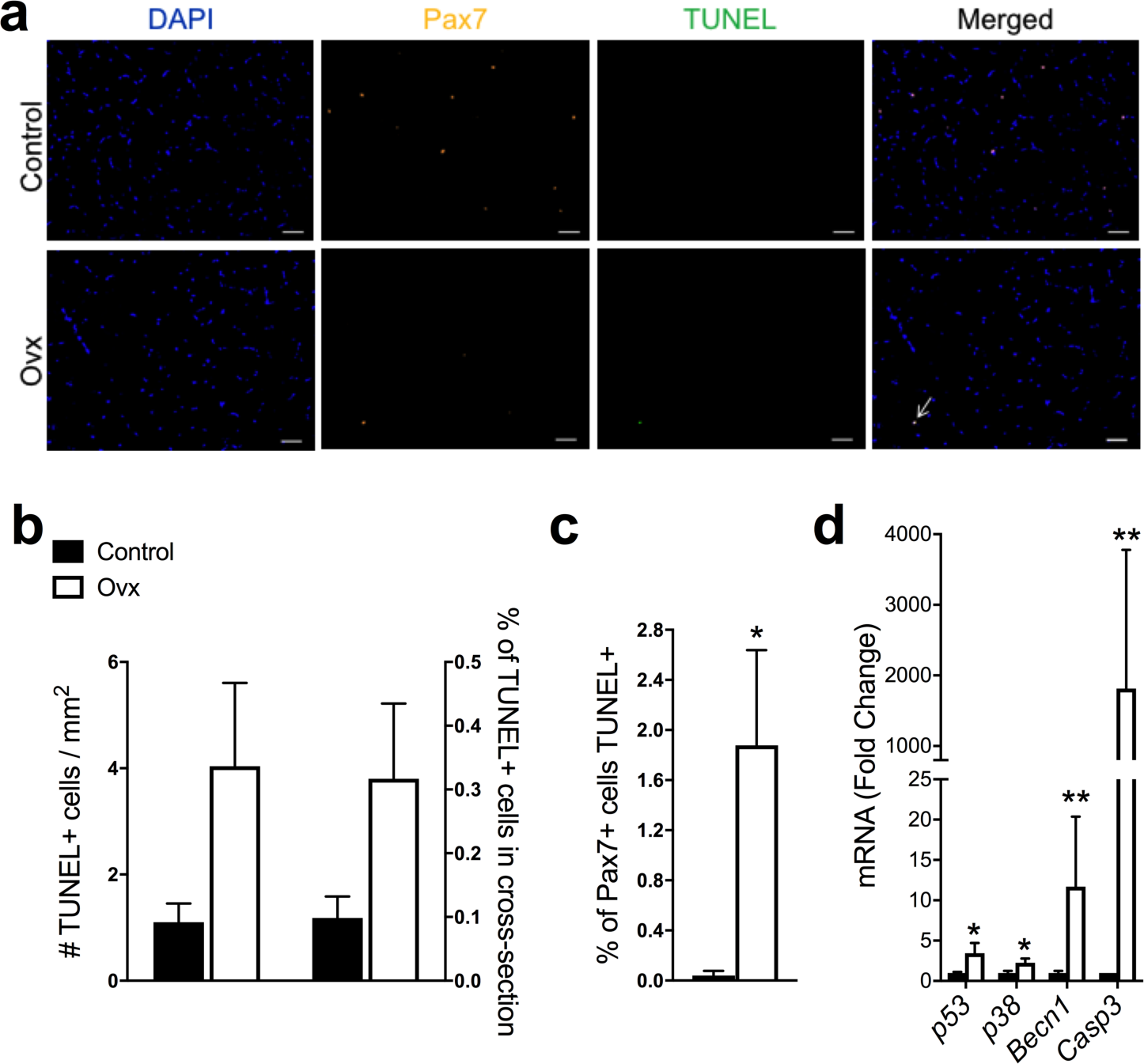
Loss of estradiol results in satellite cell apoptosis. **(a)** Representative images of DAPI stained nuclei (blue), Pax7 satellite cells (gold), TUNEL cells (green), and merged image from cross-sections of TA muscles from Control (n=4) and Ovariectomized (Ovx; n=4) mice. Arrows indicate DAPI+ Pax7+ TUNEL+triple-positive cell. Scale bars = 50 µm. **(b)** Quantification of total number of TUNEL+ cells per cross-sectional area and % TUNEL+ cells in each cross-section relative to the total # of muscle fibers. **(c)** Percent of Pax7+ cells that were also TUNEL+. **(d)** mRNA expression of apoptosis-related genes in ZsGreen+ satellite cells isolated from gastrocnemius muscles of Control (n=4) and Ovx mice (n=4). * P<0.05 and **P<0.005 by student t-tests

To determine whether any of these newly induced apoptotic cells were actually satellite cells, we counterstained samples with Pax7. Loss of estradiol resulted in 2% of the Pax7 population being apoptotic (i.e., Pax7+TUNEL+ cells), while in controls the frequency was close to zero (0.03%; Fig. 4c). To validate these results from the level of transcription, we then isolated Pax7-ZsGreen+ cells from Control and Ovx mice and evaluated the transcription of apoptosis-related genes. Estradiol deficiency resulted in 3-to 1500-fold upregulation of *p53, p38, Becn1*, and *Casp3* gene expression (Fig. 4d).

### ERα is the relevant estrogen receptor in satellite cells

There are three known estrogen receptors in skeletal muscle: estrogen receptor alpha (ERα), estrogen receptor beta (ERβ), and g-protein coupled estrogen receptor (GPER). We have recently established that the effects of estradiol on force generation by skeletal muscle are mediated by ERα (Collins et al., 2018). This result, in combination with other work showing ERα is involved in regulating muscle metabolism (Hamilton et al., 2016; Ribas et al., 2016; Ronda et al., 2013) and oxidative stress (Baltgalvis et al., 2010; Vasconsuelo et al., 2008), led us to determine 1) whether satellite cells express ERα and 2) whether this receptor was essential in mediating the activity of estradiol on satellite cell function. qRT-PCR on sorted Pax7-ZsGreen+ cells showed that satellite cells do indeed express *Esr1,* the gene encoding ERα (Fig. 5a).

**Figure 5.**
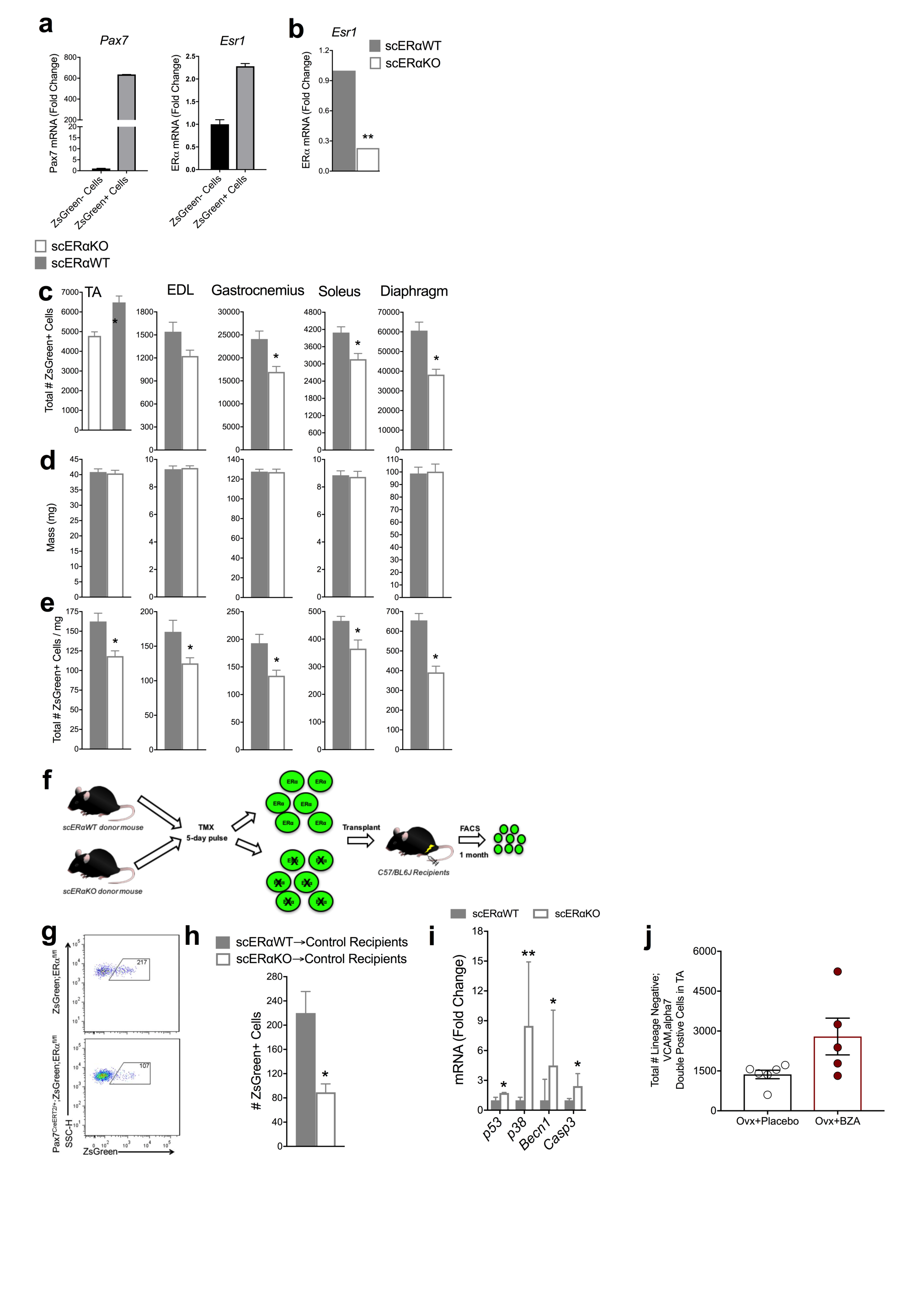
Estradiol utilizes ERα to regulate satellite cell maintenance and self-renewal. **(a)** mRNA gene expression of *Pax7* and estrogen receptor alpha (*Esr1*) in isolated ZsGreen+ satellite cells from Pax7-ZsGreen female mice (n=4). **(b)** mRNA gene expression of *Esr1* in Pax7-ZsGreen;ERα^fl/fl^ (scERαWT; n=4) and Pax7^CreERT2/+^; Pax7-ZsGreen;ERα^fl/fl^ (scERαKO; n=4) female mice. **(c)** Total number of ZsGreen+ satellite cells in five muscles of scERαWT (n=6) and scERαKO (n=6) mice. **(d)** Muscle masses. **(e)** Total number of ZsGreen+ cells normalized to muscle mass. **(f)** scERαWT and scERαKO transplantation scheme. **(g)** Representative FACS plots of ZsGreen+ donor satellite cells from scERαWT (n=3) and scERαKO (n=3) mice that were transplanted into Control recipients (n=12). **(h)** Quantification of ZsGreen+ donor satellite cells in Control recipient TA muscles following transplantation. **(i)** mRNA gene expression of *p53, p38, Becn1*, and *Casp3* in isolated ZsGreen+ satellite cells from gastrocnemius muscles of scERαWT (n=4) and scERαKO (n=4) mice. **(j)** Total number of satellite cells isolated from TA muscles of Ovx mice without (n=6) and with Bazedoxifine (Ovx+BZA; n=5) treatment (P=0.055). *P<0.05 and **P<0.005 by student t-tests

Therefore, we developed an inducible satellite cell-specific ERα knockout model (Fig. 5b) in order to specifically probe the necessity of ERα in the mechanism by which estradiol mediates effects on satellite cells. We treated Pax7^CreERT2/+^;Pax7-ZsGreen;ERα^fl/fl^ female mice (scERαKO) and Pax7^CreERT2 +/+^; Pax7-ZsGreen;ERα^fl/fl^ control sibling females (scERαWT) with tamoxifen to ablate ERα and evaluated satellite cell frequency in multiple muscle groups by FACS (Supplementary Fig. 3). Ablation of ERα in satellite cells resulted in 40-60% fewer satellite cells in all five muscles analyzed; again, both expressed in terms of absolute cell numbers and satellite cell density, i.e., normalized to muscle mass (Fig. 5c-e). We replicated these results by performing Pax7 immuno-staining of TA muscle cross-sections from scERαWT and scERαKO female mice (Supplementary Fig. 3). The magnitude of the satellite cell reductions was similar to those of estradiol deficiency (compare Fig. 5c-e with Fig. 1a-c). These results indicate mechanistically that ERα is necessary for estradiol signaling to maintain the satellite cell pool in muscles of females.

We next tested whether the loss of ERα in satellite cells would impair self-renewal using the *in vivo* transplantation assay, similar to what was measured with estradiol deficiency (Fig. 5f). 600 ZsGreen+ satellite cells from female scERαWT and scERαKO mice were transplanted into control C57/BL6J recipient mice and contribution to the satellite cell compartment was measured by FACS analysis of total recipient muscle, one-month post-transplant (Fig. 5g). Analogous to the Ovx study, we observed a dramatic reduction in satellite cell engraftment in the absence of ERα in donor satellite cells (Fig. 5h), supporting the interpretation that ERα is the relevant estrogen receptor maintaining homeostatic control of self-renewal in response to estradiol.

Finally, we evaluated the effect of ERα deletion on apoptosis. First, we counted TUNEL+ satellite cells in TA muscle cross-sections (Supplementary Fig. 3) and measured apoptosis-related transcripts in satellite cells isolated from scERαWT and scERαKO mice. In the absence of satellite cell ERα, TUNEL+ cells increased, although in this case, statistical significance was not achieved (P=0.124; Supplementary Fig. 3). However, an upregulation of apoptosis-related genes (Fig. 5i) similar to that seen with estradiol deficiency (Fig. 4) was measured.

### Bazedoxifine (BZA) acts as a selective ER agonist in satellite cells

It was not known whether the new selective estrogen receptor modulator, BZA, would be an ER agonist or antagonist in skeletal muscle. We show that Ovx mice treated with BZA rescued the detrimental effects on satellite cell number (P=0.055; Figure 5j) demonstrating that it is an agonist in muscle. Because BZA functions by binding estrogen receptors, this result further supports the concept that estradiol’s mechanism of action in satellite cells is through ERα.

### In humans, satellite cell number declines during the menopausal transition

Satellite cell number declines with age in humans, however, the majority of studies have compared muscle from old men (∼70 yr old) to young men (∼20 yr old) and the studies have been cross-sectional in design (Verdijk et al., 2012; Verdijk et al., 2014). To date, satellite cell number has not been evaluated during the menopausal transition in women, nor to our knowledge have any aging longitudinal studies of satellite cell number been conducted in humans. Here, we aimed to determine if satellite cell number declines during the peri-to post-menopausal transition in women by taking two muscle biopsies from the same women at peri- and post-menopause, and staining these for Pax7 (Fig. 6a, b). Satellite cell number declined 15% on average during the transition and trended very close to statistical significance (P=0.057; Fig. 6c), and declines were observed in 4 out of the 5 women (Fig. 5d). This reduction is striking since the peri-to post-menopause transition in these women occurred within a one-year time period and cell number declines cannot be attributed to reduced physical activity or myofiber size (Fig. 6a). We observed that not all study participants displayed the same degree of estradiol loss across the time in which longitudinal biopsies were obtained and informed by our results in mice that estradiol deficiency drove the reduction of satellite cells, we performed a Pearson correlation between serum concentration of estradiol (Fig. 6e) and satellite cells in peri- and post-menopausal samples from the same five women. The result is a strong correlation that met statistical significance (r^2^ = 0.478 P=0.023; Fig. 6f), indicating that even with the low sample size not yet allowing the acceptance of the hypothesis of a sudden satellite cell decline at the peri-post menopausal transition, the data support that changes in satellite cell number across this transition are related to changes in 17β-estradiol concentration.

**Figure 6.**
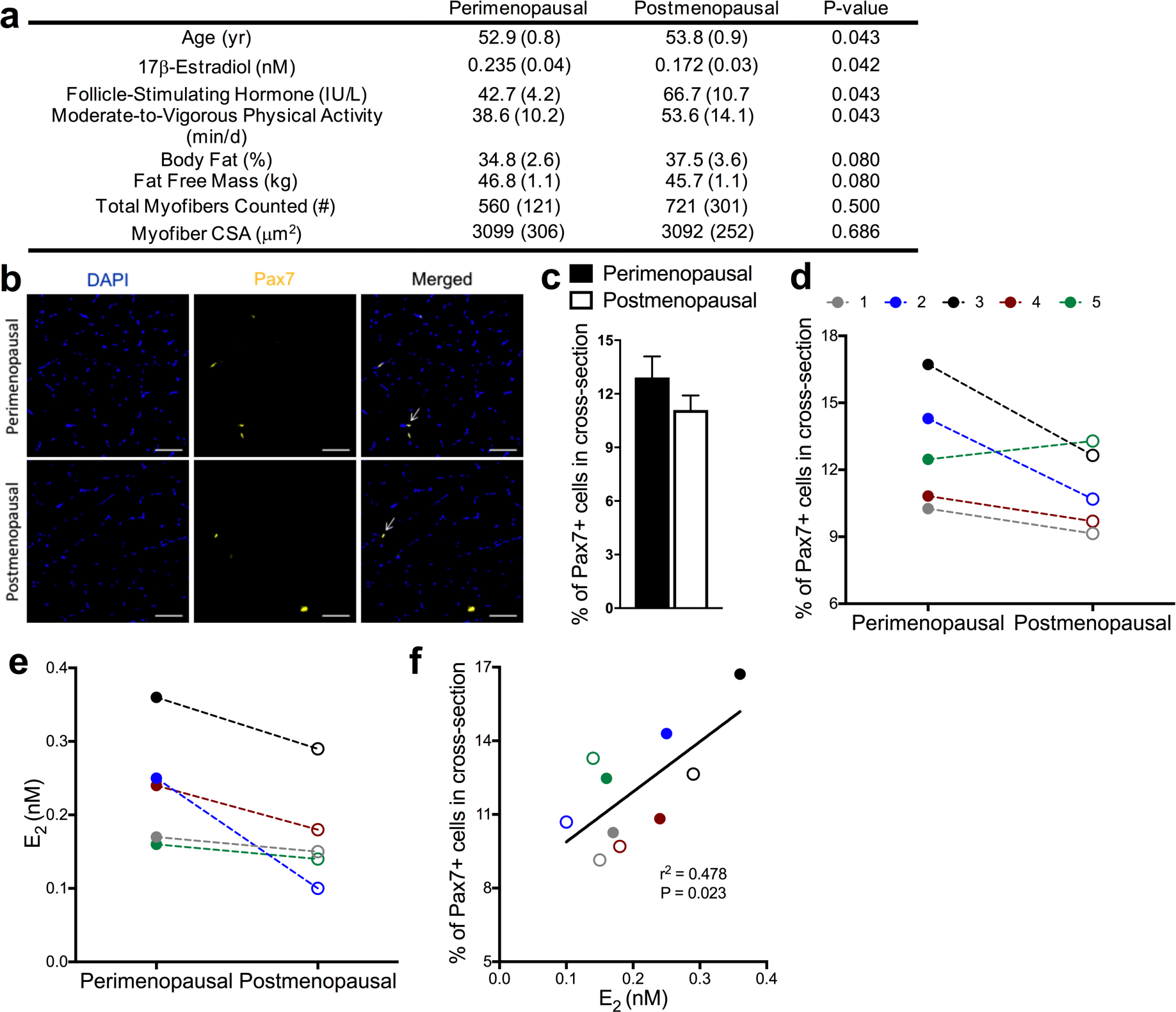
Peri-to post-menopausal transition results in decline of muscle satellite cells in humans. **(a)** Subject characteristics of women participants (n=5). **(b)** Representative images of DAPI stained nuclei (blue), Pax7 satellite cells (gold), and merged images from cross-sections of muscle biopsies from perimenopausal women and of the same women when postmenopausal status was reached. Scale bars = 50 µm. **(c)** Quantification of % of Pax7+ cells counted in each cross-section relative to the total number of fibers in muscle biopsies from Perimenopausal and Postmenopausal women (P=0.057). **(d)** Declines in the % of Pax7+ cells in muscle cross-section of 4 out of 5 individual women during the two menopausal stages. **(e)** Individual serum estradiol (E_2_) levels in women during the two menopausal stages. **(f)** Pearson correlation between E_2_ and %Pax7+ cells (P=0.023). Perimenopausal women shown in closed circles and postmenopausal women shown in open circles (def).

## DISCUSSION

The notion that age-associated deficits in skeletal muscle can be driven by extrinsic factors that differ between young and aged environments is well established by heterochronic parabiosis experiments (Brack et al., 2007; Conboy et al., 2005). Estradiol is a particularly interesting hormone in this regard as its levels are much higher in females than in males (Nelson and Bulun, 2001), levels fall precipitously in females at the menopausal transition (Baber et al., 2016), and hormone therapy has been found to have beneficial effects on skeletal muscle health in post-menopausal women (Phillips et al., 1993; Qaisar et al., 2013). To date, mechanisms of estradiol on skeletal muscle have focused primarily on effects on protein synthesis (Kamanga-Sollo et al., 2010; McClung et al., 2006), indirect anti-inflammatory effects (Le et al., 2015; MacNeil et al., 2011; McClung et al., 2007; Ribas et al., 2010), and impact on force generation (Lai et al., 2016; Moran et al., 2007). The results presented here show that estradiol is necessary for satellite cell maintenance and function in females. When estradiol is lacking in the systemic environment, the number of satellite cells is reduced by 30-50% in 4 of 5 skeletal muscles studied. The consequence of the estradiol-deficient mediated loss of satellite cells is blunted recovery of strength following injury. Strength recovery from injury is exacerbated by the second-bout of injury suggesting that satellite cell self-renewal or differentiation is impaired.

To this point, satellite cells harvested from an estradiol-rich environment failed to populate the satellite cell compartment when transplanted into a host environment that lacked estradiol, but when satellite cells from the pool remaining in a hormone-deficient environment were transplanted into a host environment with estradiol, engraftment, self-renewal, and differentiation were robust. Moreover, we measured a similar decline in satellite cell pool size when ERα is deleted specifically from satellite cells in steady state muscle. These results strongly suggest a cell-autonomous role for estradiol signaling through ERα in the satellite cell itself, the impairment which leads to a reduction in satellite cell pool size.

In principal, a reduction in stem cell pool size can be explained by three possible mechanisms: differentiation, death, or emigration of cells from the tissue. We find no evidence of increased differentiation in the estradiol-deficient environment. Rather, the opposite appears to hold: when lineage-marked cells were transplanted there was greater contribution to differentiated muscle fibers in the estradiol-replete environment than in the deficient environment. Given that an impairment in differentiation would tend to increase the number of stem cells, not decrease it, we feel that the lack of contribution to differentiated muscle in this assay is not due to a differentiation blockade. Rather, because we find a subpopulation of Pax7+ cells that are TUNEL+, together with a very strong apoptotic transcriptional response, it is most likely that Pax7+ cells are being lost to apoptosis. Estradiol is known for anti-apoptotic effects in some other contexts, for example studies on C2C12 cells provide evidence that estradiol can be protective from H_2_O_2_-induced apoptosis through modulating P53 and FOXO transcription factors and their downstream target genes (Boland et al., 2008; La Colla et al., 2016). Concerning the third possibility, although our studies do not formally rule out emigration of satellite cells from estradiol-deficient muscle, the scenario seems unlikely due to the fact that satellite cells do not repopulate via the circulation into muscles that have been completely depleted by freeze injury (although they can migrate into adjacent muscles if connective tissues are disrupted: Schultz et al., 1986).

Hormone therapy is controversial because of the small but detectable increase in breast cancer risk (Kim et al., 2018). The etiology of this disease is thought to be related to estradiol stimulation or mutations in the gene ERα (Dall et al., 2018; Gelsomino et al., 2018; Severson et al., 2018), which is known to exert proliferation of the normal breast tissue (Dall et al., 2018). However, pharmacological progress has been made and there now exists FDA approved selective estrogen receptor modulators (SERMs) (Börjesson et al., 2016). Because SERMs function by inducing conformational changes in ER resulting in tissue-specific agonistic or antagonistic effects, largely depending on which cofactors are present, it was not clear whether the SERM, BZA, would present as an agonist or antagonist in skeletal muscle (Beck et al., 2015; Börjesson et al., 2016). Our results suggest that BZA is an ER agonist in skeletal muscle, or at least in muscle stem cells, as it restored the satellite cell pool in estradiol-deficient mice. This result along with our finding that ERα is the relevant receptor whose signaling is necessary to maintain the satellite cell pool, supports the use of ERα-selective ligands for therapeutic application in menopause-associated muscle loss.

Notably, we also present preliminary data from the first longitudinal biopsy study of aging skeletal muscle. This study tracks muscle in women and employs variable biopsy dates. The first biopsy is at peri-menopause, and the second at post-menopause, with timing being unique to each participant and determined by changes in their estradiol and follicle stimulating hormone levels. The time between first and second biopsy was short (∼ 1 year). Accordingly, although we observed a reduction in mean satellite cell number from first biopsy to second, we observed a much greater significance in the correlative analysis between changes in estradiol levels with satellite numbers, this comparison actually being the most relevant to the current study.

In summary, our results demonstrate that estradiol is a necessary factor regulating *in vivo* satellite cell function in female rodents and is supported by similar results in humans. The estradiol-ERα axis contains potential therapeutic targets to mitigate skeletal muscle deficits observed with aging in women. Our study provides scientific basis for future studies to examine ER agonists, such as BZA, more thoroughly to probe for additional benefits toward overall skeletal muscle health in postmenopausal women. Finally, these collective results highlight the importance of considering biological sex and sex hormones when studying skeletal muscle, particularly regarding age-related deficits in muscle.

## ACKNOWLEDGEMENTS

We thank Angus Lindsay, Vineesha Kollipara, Tara Mader, Mayank Verma, Alessandro Magli, Yi Ren and Olivia Recht for technical assistance and Atsushi Asakura and Ken Korach/Andrea Hevener for the Pax7^CreERT2/+^;R26R^tdTomato^ mice and ERα floxed mice, respectively. Bazedoxifene was a gift from Pfizer.

## GRANT SUPPORT

Our research has been supported by the National Institutes of Health Grants R01-AG031743 (DAL), R01-AR055685 (MK), and T32-AR007612 (BCC, RWA, CWB, and CAC), the Muscular Dystrophy Association grant MDA351022 (MK), a grant from the American Diabetes Association BS-1-15-170 (EES), a grant from the Office of the Vice President for Research, University of Minnesota (DAL), and grants from the Academy of Finland 275323 (VK) and 309504 (EKL). BCC was also supported by a University of Minnesota Interdisciplinary Doctoral Fellowship.

## AUTHOR CONTRIBUTIONS

BCC: Study design, data collection, analysis, interpretation, manuscript writing

RWA: Study design, data collection, analysis, interpretation

AAL: Data collection, analysis, interpretation

CWB: Data collection, analysis, interpretation

CAC: Data collection, analysis

NLN: Data collection, analysis

HKJ: Human experiments and analysis

EKL: Human study design, experiments and analysis, financial support

SS: Human study design and experiments and analysis

VK: Human study design and experiments, financial support

EES: Data collection, analysis

MK: Study design, data interpretation, manuscript writing, financial support

DAL: Conception and design, data interpretation, manuscript writing, financial support

## DISCLOSURE OF POTENTIAL CONFLICTS OF INTEREST

The authors indicate no potential conflicts of interest.

## STAR METHODS

### Mice

Animal experiments in this study were performed in accordance with protocols approved by the Institutional Animal Care and Use Committee at the University of Minnesota. All experiments were conducted on female mice when they were young adults (3-4 mo of age). Satellite cells were harvested from C57/BL6 (Purchased from Jackson Laboratories, 000664) and Pax7-ZsGreen mice that were either control or ovariectomized (bilateral removal of the ovaries) and sacrificed 2, 3.5, or 7 mo post-surgery (n=5 for each group). ERα^fl/fl^ mice were ovariectomized (Ovx) and half received a 0.18 mg 60-day slow-release 17β-estradiol (Ovx+E_2_) pellet via trochar implantation (n=3-4) (Moran et al., 2007). For strength experiments, C57/BL6 mice were ovariectomized and half received placebo pellets (Ovx+placebo) and half received 0.18 mg 60-day slow-release 17β-estradiol pellets (Ovx+E_2_) (n=6-8). For Bazedoxifine experiments, C57/BL6 mice were ovariectomized and half received placebo pellets (Ovx+placebo) and half received 1.44 mg Bazedoxifine (Ovx+BZA) pellets (n=5-6) (Andersson Annica 2016). For transplant experiments, female Pax7-ZsGreen and Pax7^CreERT2/+^;R26R^tdTomato^ mice were donors and were age-matched to female C57/BL6 recipient mice (n=6-15 for each experiment); a subset of recipients were ovariectomized. Pax7^CreERT2/+^;Pax7-ZsGreen;ERα^fl/fl^ (scERαKO) and Pax7-ZsGreen;ERα^fl/fl^ (scERαWT) were generated in-house. Female scERαWT and scERαKO mice were treated with tamoxifen for 5 d consecutively (Keefe et al., 2015; Murphy et al., 2011). Two mo after tamoxifen treatment, mice were used for satellite cell harvests (n=6) and transplantation (n=6). For all experiments, the estrous cycle was tracked for 3-5 d consecutively via vaginal cytology to confirm mice had normal estrous cycles or had ceased cycling for those that were Ovx (Nelson et al., 1982). Uterine mass was measured at the time of sacrifice as a second verification of successful ovariectomy surgery or estradiol treatment (Wood et al., 2007). Uterine mass across all experiments for control, Ovx, and Ovx+E_2_ mice averaged (SEM): 101.7 (5.1), 13.5 (0.7), and 207.9 (27.0) mg, respectively.

### Human Participants

Muscle samples from the Estrogenic Regulation of Muscle Apoptosis (ERMA) Study were analyzed for satellite cells. The ERMA study is a population-based longitudinal cohort study comprised of women 47 to 55 years of age living in the city of Jyväskylä and the neighboring municipalities in Finland (Kovanen V, Submitted). Participants were assigned to menopausal groups following STRAW +10 guidelines (Harlow et al., 2012). Systemic follicle stimulating hormone (FSH) and 17β-estradiol levels were immunoassayed by IMMULITE^®^ 2000 XPi System (Siemens Healthcare Diagnostics, UK) from the fasting blood samples taken from the antecubital vein in a supine position between 7:00 and 10:00 AM. Percentage of body fat and lean body mass was assessed by a multifrequency bioelectrical impedance analyzer (InBody^™^ 720; Biospace, Seoul, Korea) after overnight fasting with the participant wearing only undergarments. Physical activity was assessed with ActiGraph accelerometers (Pensacola, Florida, USA) as in Laakkonen et al(Laakkonen et al., 2017). Briefly, participants were instructed to wear monitors on the right hip during waking hours for seven consecutive days. Moderate to vigorous intensity physical activity (MVPA) was defined by using tri-axial vector magnitude cut-point over 2690 counts per minute (cpm) to define the cut-point for at least moderate intensity level. There were small intra-individual differences in wearing times of the accelerometer (mean wearing time 15.2±0.3 hours/day across 6.8±0.2 days). Therefore, the MVPA time (min/day) was normalized by wearing time to correspond daily 16 hour waking time. The Ethics Committee of the Central Finland Health Care District approved the ERMA Study in 2014 (K-S shp Dnro U/2014). An informed consent was given by each participant at the laboratory before any sampling or measurement was done. Altogether, the study protocol followed good clinical and scientific practice and the Declaration of Helsinki.

### Mouse Satellite Cell Isolation

Isolation of satellite cells from single skeletal muscles (e.g., gastrocnemius) was performed as described previously (Arpke and Kyba, 2016). Muscles were carefully dissected and chopped in parallel with muscle fibers using razor blade and forceps to separate the fibers. Muscles were incubated shaking for 75 min in 0.2% collagenase type II (17101-015, Gibco, Grand Island, NY) in high glucose Dulbecco’s modified Eagle’s medium (DMEM) without phenol red containing 4.00 mM L-glutamine, 4,500 mg/L glucose, and sodium pyruvate (SH30284.01, Hyclone, Logan, UT) supplemented with 1% Pen/Strep (15140122, Gibco) at 37oC. Samples were washed with Rinsing Solution (F-10+), Ham’s/F-10 medium (SH30025.01, HyClone) supplemented with 10% Horse serum, 1% HEPES buffer solution (15630080, Gibco) and 1% Pen/Strep (Gibco) and centrifuged at 1500 rpm × 5min at 4°C. Samples were washed and centrifuged a second time. Samples were pulled into a sheared Pasteur pipette, centrifuged and washed again. Following aspiration, samples were resuspended in F-10+ with collagenase collagenase type II and dispase (17105-041, Gibco), vortexed and incubated shaking at 37°C for 30 min. Samples were vortexed again, drawn and released into a 3 mL syringe with 16-gauge needle four times then with an 18-gauge needle four times and passed through a 40-μm cell strainer (Falcon, Hanover Park, IL). 3 mL of F-10+ was added to each sample and centrifuged at 1500 rpm × 5min 4°C. Following aspiration, samples were resuspended in FACS staining medium (2% FBS in PBS). Bulk isolation (hindlimb muscles excluding soleus, triceps muscles, and psoas muscles) of satellite cells was performed similarly and as described previously (Arpke et al., 2013).

### FACS Analysis and Cell Sorting

Muscle samples were stained using an antibody mixture of PE-Cy7 rat anti-mouse CD31 (clone 390), PE-Cy7 rat anti-mouse CD45 (clone 30-F11), Biotin rat anti-mouse CD106 (clone 429(MVCAM.A)) and PE Streptavidin from BD Biosciences (San Diego, CA); and Itga7 647 (clone R2F2) from AbLab (Vancouver, B.C., Canada). Antibody cocktail was added to samples and incubated on ice for 30 min. Samples were washed and resuspended with FACS staining media containing propidium iodide for FACS analysis on the FACSAriaII SORP (BD Biosciences, San Diego, CA). Total satellite cells (lineage negative; VCAM,alpha7 double positive cells or ZsGreen+) (Supplementary Fig. 1a,b) were analyzed while draining the entire sample from each skeletal muscle sample. For transplanted tibialis anterior (TA) muscles, the number of donor (ZsGreen + or tdTOM+) satellite cells were examined as previously described (Arpke et al., 2013).

### Barium Chloride Injury and *In Vivo* Muscle Torque Measurement

As previously described (Baltgalvis et al., 2009; Baumann et al., 2014; Lowe et al., 1995), contractile function of the left anterior crural muscles was measured *in vivo* immediately before the injuries, as well as 7, 14, and 21 days after each injury. Briefly, anesthetized mice (1.25% isoflurane and 125 mL O_2_ per minute) were placed on a temperature-controlled platform to maintain core body temperature between 35 and 37 °C. The left knee was clamped and the left foot was secured to an aluminum “shoe” that is attached to the shaft of an Aurora Scientific 300B servomotor (Aurora Scientific, ON, Canada). Sterilized platinum needle electrodes were inserted through the skin for stimulation of the left common peroneal nerve. Stimulation voltage and needle electrode placement were optimized with 5–15 isometric contractions (200 ms train of 0.1 ms pulses at 200 Hz). Two minutes following optimization, contractile function of the anterior crural muscles was assessed by measuring isometric torque as a function of stimulation frequency (20–300 Hz), with the highest recorded torque defined as maximal isometric torque. Following pre-injury and 21 day torque measurements, barium chloride (1.2% in sterile demineralized water) (Ricca Chemical Company, Arlington, TX) was injected into the tibialis anterior muscle of each mouse with a Hamilton syringe similar to Murphy et al (Murphy et al., 2014). Functional torque measurements were performed on the same groups of mice at each time-point post-injury, and expressed relative to pre-injury maximal isometric torque.

### Cardiotoxin Injury and Transplantation

Transplant recipient mice were anesthetized with 150 mg/kg ketamine plus 10 mg/kg xylazine and both hind limbs were subjected to a 900 cGy dose of irradiation using an RS 2000 Biological Research Irradiator (Rad Source Technologies, Inc., Suwanee, GA). Lead shields limited exposure to the hind limbs only. 24 h following irradiation, 15 μL of cardiotoxin (10 μM in PBS, Sigma-Aldrich, Saint Louis, MO) was injected into both TA muscles of each mouse with a Hamilton syringe. 24 h following cardiotoxin injection, 600 ZsGreen+ cells were resuspended into 10 μL of sterile saline and injected into both TA’s. Both TA’s were harvested 1 mo post-transplantation and prepared for FACS analysis as described above. When tdTom+ cells were transplanted, one TA was harvested and prepared for FACS analysis and the contralateral TA was harvested and prepared for sectioning and staining.

### Immunofluorescence Microscopy in Mouse Samples

TA muscles were removed and placed in OCT compound, frozen in 2-methylbutane (Sigma-Aldrich) cooled by liquid nitrogen and stored at -80°C until use. For visualization of satellite cells and apoptotic cells in skeletal muscle, Pax7 and terminal deoxynucleotidyl transferase dUTP nick end labeling (TUNEL) staining was performed on 7 μM cryosections (CM 1850, Leica Microsystems, Buffalo Grove, IL). TUNEL was detected using In Situ Cell Death, Fluorescein Detection Kit according to manufacturer’s instructions (11684795910, Roche Diagnostics, Mannheim, Germany). Following TUNEL, Pax7 staining was completed as previously reported (Keefe et al., 2015; Murphy et al., 2011). In brief, slides were incubated in citrate buffer for 10 min and then antigen retrieval was performed by microwaving at 20 sec intervals for a total of 3 min in citrate buffer. Slides were washed two times in 1x PBS for 2 min each and blocked in 3% bovine serum albumin (BSA) for 1 h at room temperature. Slides were incubated with anti-pax7 mouse IgG1 primary antibody (PAX7, Developmental Hybridoma Bank at Iowa University, 1:20 in 3% BSA) overnight at 4°C. Slides were incubated with goat anti-mouse biotin-conjugated secondary antibody (115-065-205, Jackson Immuno Research Laboratories Inc, West Grove, PA, 1:1000 in 3% BSA) for 1 h at room temperature.

Visualization of the primary antibody was performed using the Vectastain ABC kit (PK-6100, Vector Laboratories, Burlingame, CA) and Tyramide Signal Amplification (TSA) Plus Cyanine 3 kit (NEL744, PerkinElmer, Waltham, MA, 1:50 in diluent buffer). Slides were washed again and then prolong gold antifade mountant with DAPI (P36931, Life technologies, Grand Island, NY) was applied and coverslip added.

For determination of the cross-sectional area of the TA muscle and to determine the myofiber border, adjacent 7 μm sections of each mouse sample were cryo-sectioned. Samples were fixed in 4% PFA for 10 min, washed with 1x PBS and blocked with 3% BSA in 1x PBS for 60 min at room temperature. Sections were incubated overnight at 4°C with primary antibody anti-laminin (L9393; Sigma-Aldrich, 1:250) diluted in 1% BSA in 1x PBS. Slides were washed with 1x PBS and secondary antibody Alexa 488 goat anti-rabbit (A11034; Life Technologies, 1:500) was applied for 1 h at room temperature. Slides were washed and then mounted with Prolong Gold Anti-fade mountant media (Life Technologies).

For visualization of fiber engraftment following transplantation, 10 μM sections of TA muscles were cryosectioned. Samples were fixed in 2% paraformaldehyde (PFA) for 5 min, washed with 0.01% triton in 1x PBS, permeabilized with 0.2% triton for 5 min, and then blocked with 1% BSA in wash buffer for 30 min at room temperature. Sections were incubated overnight at 4°C with primary antibodies diluted in 1% BSA in wash buffer simultaneously; rabbit polyclonal anti-RFP (600-401-379, Rockland Immunochemicals Inc, Pottstown, PA, 1:200) and mouse monoclonal anti-laminin (Clone LAM-89; Sigma-Aldrich, 1:250). Slides were washed with wash buffer and secondary antibodies were applied for 1 h at room temperature simultaneously, Alexa 555 goat anti-mouse and Alexa 480 goat anti-rabbit IgG (A-21422 and A-11034, 1:500; Life Technologies). Slides were washed and then mounted with prolong gold anti-fade mountant media (P10144, Life Technologies).

### Immunofluorescence Microscopy in Human Samples

Needle muscle biopsies were obtained under local anaesthesia from vastus lateralis of five women when they were perimenopausal and then again after being classified as postmenopausal. Visible connective, adipose tissue and blood was removed before the biopsy sample was embedded in Tissue Tek^™^ compound and frozen in 2-methylbutane (Sigma-Aldrich) cooled in liquid nitrogen and stored at -150°C until use. Pax7 positive cells were identified on 10 μm cryosections (CM 3000, Leica Instruments). In brief, slides were incubated in citrate buffer for 10 min and then antigen retrieval was performed by microwaving for 4 min in citrate buffer. Slides cooled for 5 min and were washed with 1x PBS. Blocking was done in 2% TNB-buffer (Blocking reagent, FP1020, PerkinElmer) for 1 h at room temperature. Slides were incubated with anti-pax7 mouse IgG1 primary antibody (PAX7, Developmental Hybridoma Bank at Iowa University, 1:25 in 2% TNB) overnight at 4°C. Slides were incubated with goat anti-mouse biotin-conjugated secondary antibody (Jackson Immuno Research Laboratories Inc, 1:1000 in 2% TNB) for 1 h at room temperature. Visualization of the primary antibody was performed using the Vectastain ABC kit (Vector Laboratories) and Tyramide Signal Amplification (TSA) Plus Cyanine 3 kit (PerkinElmer, 1:50 in diluent buffer). Slides were washed again and then Prolong Gold Antifade mountant with DAPI (Life technologies) was applied.

For determination of the cross-sectional area of myofibers, a serial 10 μm section of each biopsy was analyzed. To do this, sections were fixed in 4% PFA for 10 min, washed with 1x PBS and blocked with 3% BSA in 1x PBS for 60 min at room temperature. Sections were incubated overnight at 4°C with rabbit anti-laminin antibody (Sigma-Aldrich, 1:250) diluted in 1% BSA in 1x PBS. Slides were washed with 1x PBS and secondary antibody Alexa 488 goat anti-rabbit was applied for 1 h at room temperature (Life Technologies, 1:500). Slides were washed and then mounted with Prolong Gold Anti-fade mountant media (Life Technologies).

### Image Processing and Analysis

All images were processed and analyzed in a blinded manner with samples being de-identified as to treatment or group. Mouse muscle samples were examined and imaged using a Leica DM5500B microscope (Leica Microsystems) at 5x-20x magnification. Images were stitched using the automated tile-scan tool to construct an image of the entire cross-section of the TA muscle. Muscle cross-sectional area (CSA) in millimeters squared was measured by tracing the muscle section border, stained with laminin, in Image J. Satellite cells were identified by DAPI+ and Pax7+ cell residing along the myofiber border. Myonuclei undergoing apoptosis were identified as DAPI+ and TUNEL+ cells that were within the myofiber. Satellite cells undergoing apoptosis were identified as Pax7+, TUNEL+ and DAPI+ residing along the myofiber border. DAPI+ Pax7+ TUNEL+ cells were normalized to muscle cross-sectional area and percent of cells in each cross-section relative to total number of myofibers. For transplantation experiments, donor cells were quantified by counting RFP+ (Red) fibers using the Image J software package (NIH, Bethesda, MD, USA).

Human muscle samples were examined and imaged using a Zeiss LSM 700 confocal microscope (Carl Zeiss MicroImaging, Jena, Germany) at 10x magnification. Images were stitched using Image J software package. The number and cross-sectional area of myofibers per muscle were counted and measured using laminin staining to delineate each myofiber border in serial sections. Satellite cells were identified as DAPI+ and Pax7+ cell residing along the myofiber border. The percent of Pax7+ cells are expressed relative to myofibers counted in each biopsy.

### qRT-PCR

RNA from freshly FACS-isolated satellite cells was isolated using direct-Zol MicroPrep kit according to manufacturer’s instructions. cDNA was synthesized from 10 ng RNA according to directions in SUPERVILO cDNA Synthesis Kit (11756050, ThermoFisher, Waltham, MA). Relative quantitation of the ERα (*Esr1*), *p53, p38*, Beclin-1 (*Becn1*) and caspase-3 (*Casp3*) transcripts were determined using TaqMan probes (ThermoFisher) for ESR1 (Mm00433149_m1), Trp53 (Mm01731290_g1), Mapk14 (Mm01301009_m1), BECN1 (Mm00477631_m1), CASP3 (Mm01195085_m1), and house-keeping gene gapdh (Mm99999915_g1).

### Statistical Analysis

Data are presented as means ± SE. Two-way analysis of variance (ANOVA) were used to determine differences among time and treatment. If an interaction was significant, Holm-Sidak post-hoc tests were used. Other mouse data were analyzed with independent *t*-tests for determining differences between groups. Human data were analyzed with Wilcoxon Signed Rank test and independent *t*-tests to determine differences between peri- and post-menopause. Analyses were conducted using SigmaPlot (version 12.5, Systat Software, Inc) for mouse data and using SPSS (version 24, IBM Corporation) for human data.

## KEY RESOURCES TABLE

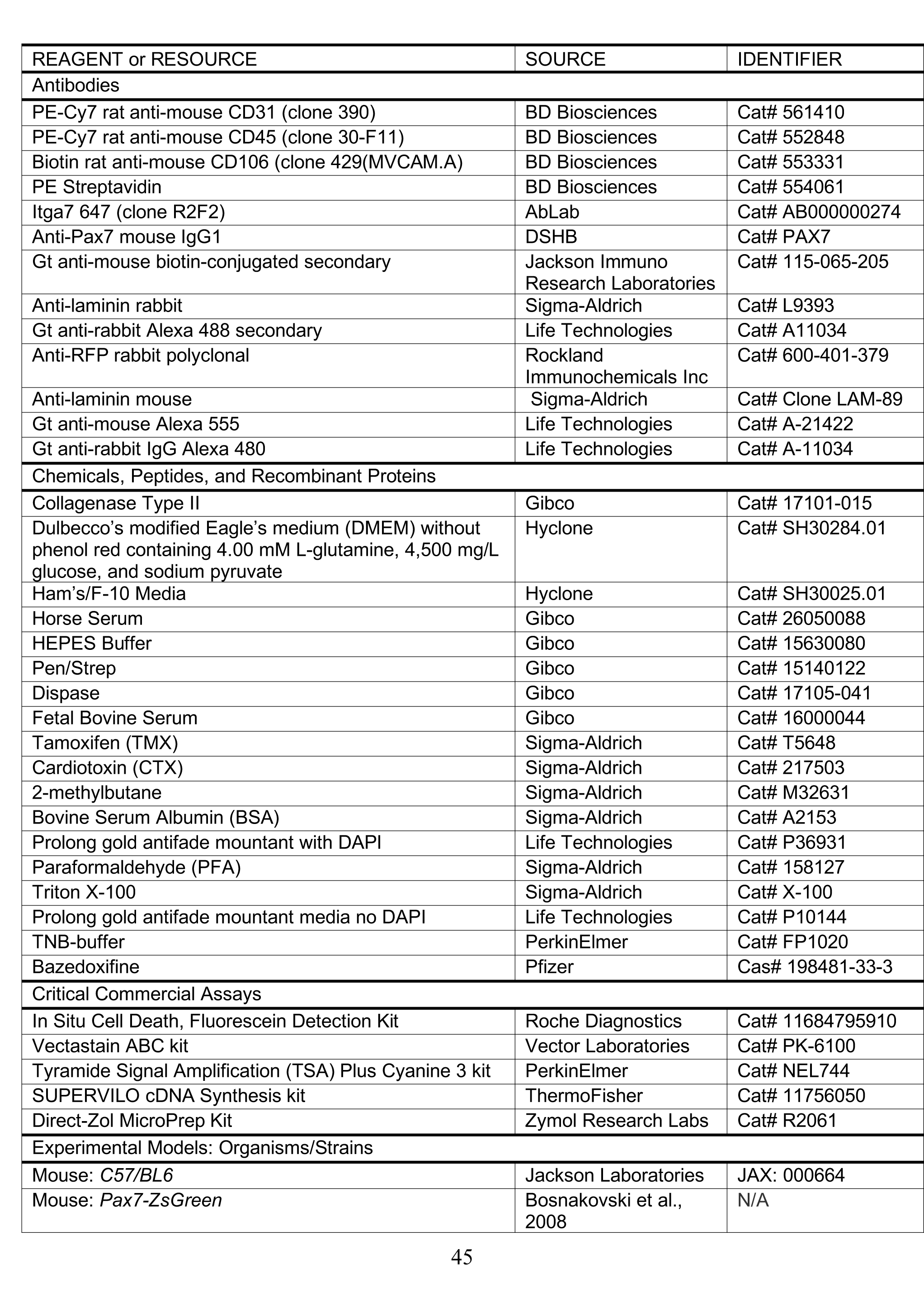
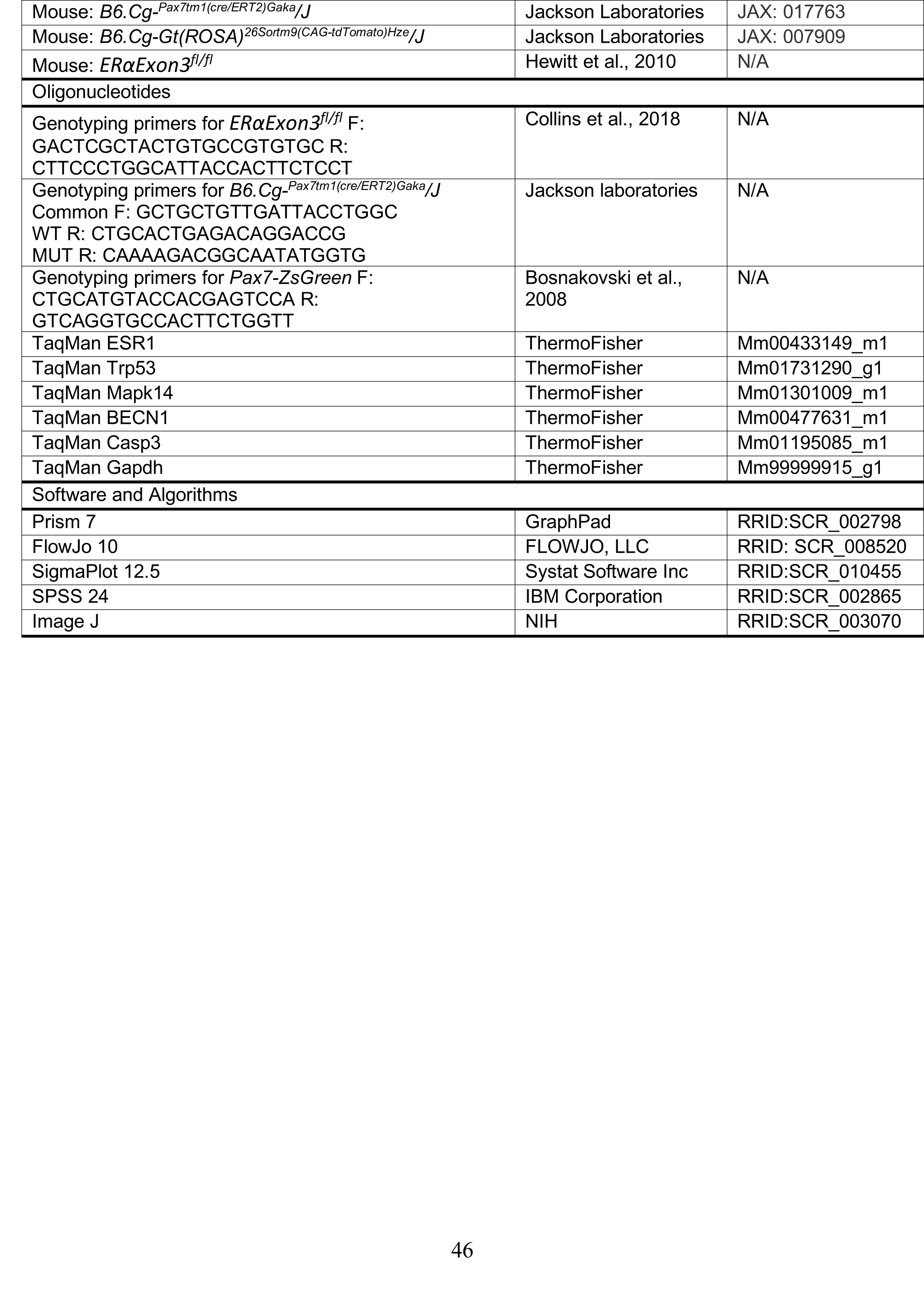

### SUPPLEMENTARY INFORMATION

**Supplementary Figure S1.**
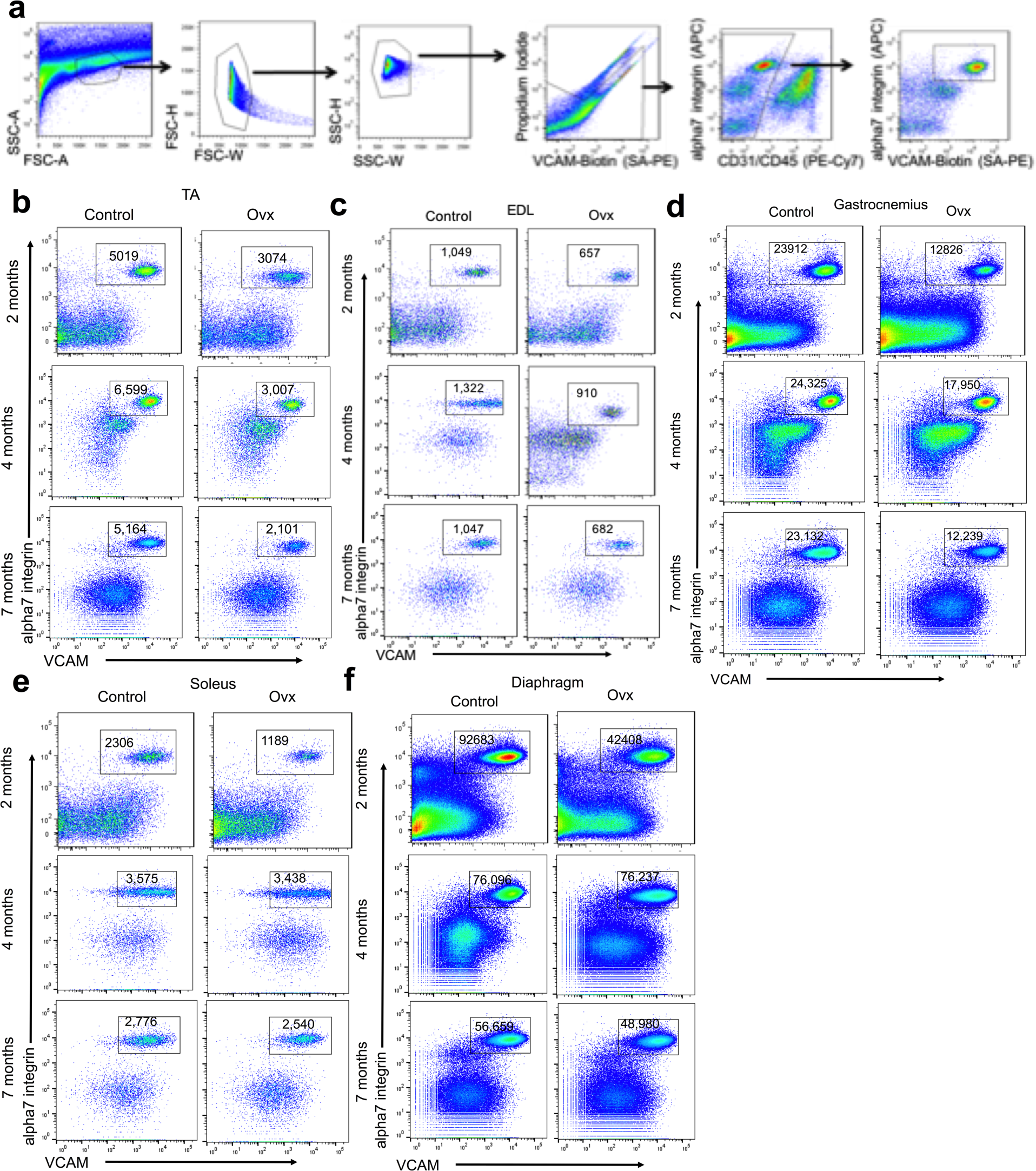
Satellite cells identified by surface marker staining **(a)** Gating scheme for lineage negative;VCAM,alpha7 double positive cells. Skeletal muscles were enzymatically digested and gated based on forward/side scatter (plots 1-3) and for live cells (propidium iodide negativity – plot 4). These cells were then gated for lineage negative cells CD31/CD45 (plot 5) and then selected for VCAM,alpha7 double positivity (plot 6). FACS plots for the quantification of the total number of satellite cells in different muscles from control and ovariectomized (Ovx) mice **(b)** Tibialis anterior (TA), **(c)** Extensor digitorum longus (EDL), **(d)** Soleus, **(e)** Gastrocnemius, **(f)** Diaphragm

**Supplementary Figure S2.**
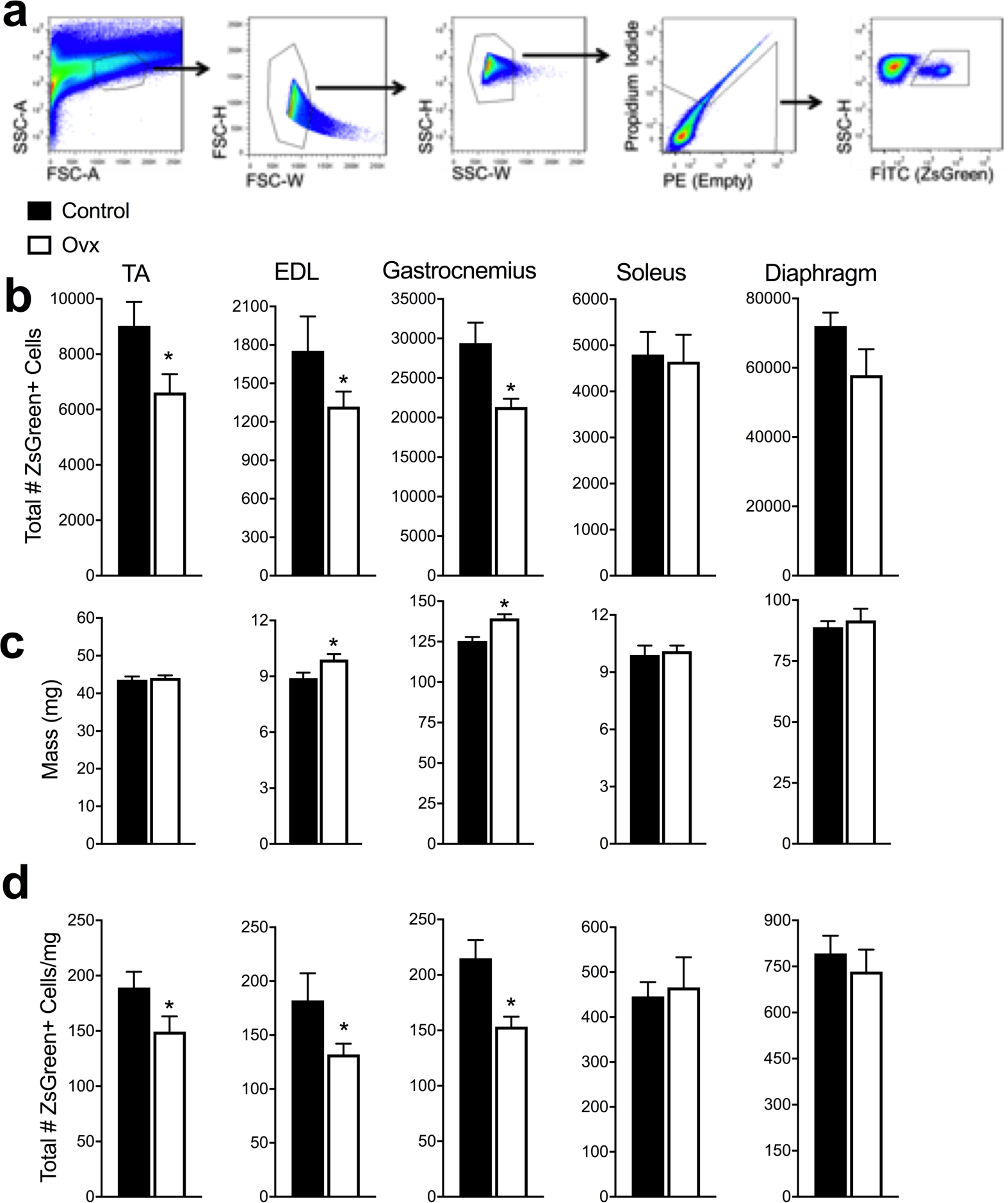
Lack of estradiol results in lower satellite cell numbers in Pax7-ZsGreen female mice. Quantification of **(a)** Total number of ZsGreen+ cells **(b)** Muscle mass **(c)** Total number of ZsGreen+ cells normalized to muscle mass in control and ovariectomized (Ovx) Pax7-ZsGreen female mice. **(d)** Gating scheme for ZsGreen+ cells. Cells were gated based on forward/side scatter (plots 1-3) and then for propidium iodide negativity (plot 4). These cells were then selected for ZsGreen positive cells SSC-H X FITC (ZsGreen).* P<0.05 by student t-tests.

**Supplementary Figure S3.**
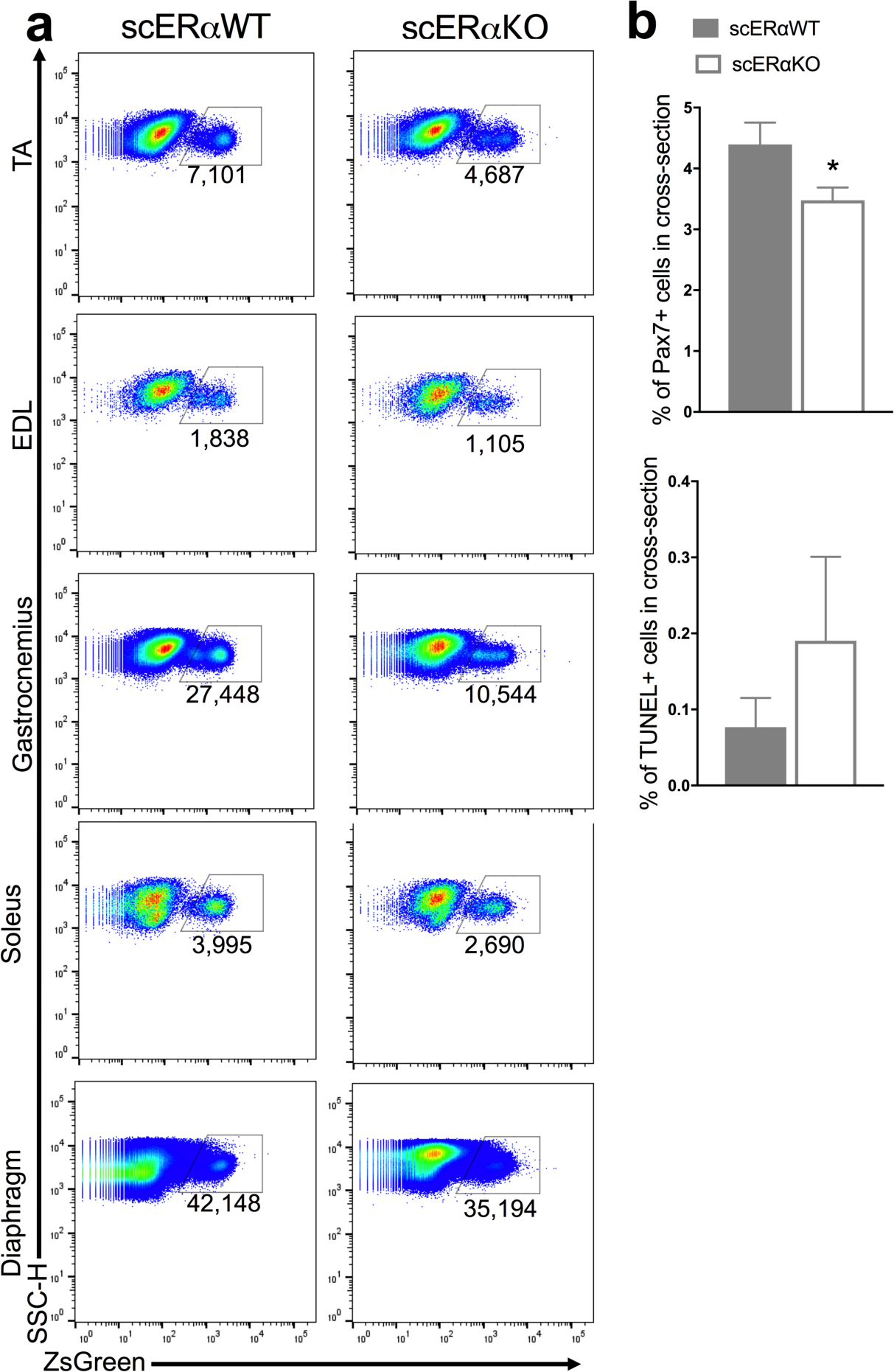
FACS plots and IHC for the quantification of the total number of satellite cells in different muscles from ZsGreen;ERα^fl/fl^ (scERαWT) (n=6) and Pax7^CreERT2/+^; ZsGreen;ERα^fl/fl^ (scERαKO) (n=6) female mice. **(a)** Tibialis anterior (TA), extensor digitorum longus (EDL), gastrocnemius, soleus, and diaphragm **(b)** % Pax7+ cells per TA muscle cross-section relative to the total # of muscle fibers. Quantification of % TUNEL+ cells in per TA muscle cross-section relative to the total # of muscle fibers.

## REFERENCES

Andersson Annica, B.A.I., Nurkkala-Karlsson Merja, Stubelius Alexandra, Grahnemo Louise, Ohlsson Claes, Carlsten Hans, Islander Ulrika (2016). Suppression of Experimental Arthritis and Associated Bone Loss by a Tissue-Selective Estrogen Complex. Endocrinology 157, 1013–1020.

Arpke, R.W., Darabi, R., Mader, T.L., Zhang, Y., Toyama, A., Lonetree, C.L., Nash, N., Lowe, D.A., Perlingeiro, R.C., and Kyba, M. (2013). A new immuno-, dystrophin-deficient model, the NSG-mdx(4Cv) mouse, provides evidence for functional improvement following allogeneic satellite cell transplantation. Stem cells (Dayton, Ohio) 31, 1611–1620.

Arpke, R.W., and Kyba, M. (2016). Flow Cytometry and Transplantation-Based Quantitative Assays for Satellite Cell Self-Renewal and Differentiation. Methods Mol Biol 1460, 163–179.

Baber, R.J., Panay, N., and Fenton, A. (2016). 2016 IMS Recommendations on women’s midlife health and menopause hormone therapy. Climacteric 19, 109–150.

Baltgalvis, K.A., Call, J.A., Nikas, J.B., and Lowe, D.A. (2009). The effects of prednisolone on skeletal muscle contractility in mdx mice. Muscle & nerve 40, 443–454.

Baltgalvis, K.A., Greising, S.M., Warren, G.L., and Lowe, D.A. (2010). Estrogen regulates estrogen receptors and antioxidant gene expression in mouse skeletal muscle. PLoS One 5, e10164.

Baumann, C.W., Rogers, R.G., Gahlot, N., and Ingalls, C.P. (2014). Eccentric contractions disrupt FKBP12 content in mouse skeletal muscle. Physiological Reports 2, e12081.

Beck, T.J., Fuerst, T., Gaither, K.W., Sutradhar, S., Levine, A.B., Hines, T., Yu, C.-R., Williams, R., Mirkin, S., and Chines, A.A. (2015). The effects of bazedoxifene on bone structural strength evaluated by hip structure analysis. Bone 77, 115–119.

Bernet, J.D., Doles, J.D., Hall, J.K., Tanaka, K.K., Carter, T.A., and Olwin, B.B. (2014). p38 MAPK signaling underlies a cell-autonomous loss of stem cell self-renewal in skeletal muscle of aged mice. Nature Medicine 20, 265–273.

Boland, R., Vasconsuelo, A., Milanesi, L., Ronda, A.C., and de Boland, A.R. (2008). 17beta-estradiol signaling in skeletal muscle cells and its relationship to apoptosis. Steroids 73, 859–863.

Börjesson, A.E., Farman, H.H., Movérare-Skrtic, S., Engdahl, C., Antal, M.C., Koskela, A., Tuukkanen, J., Carlsten, H., Krust, A., Chambon, P., et al. (2016). SERMs have substance-specific effects on bone, and these effects are mediated via ERαAF-1 in female mice. American Journal of Physiology-Endocrinology and Metabolism 310, E912–E918.

Bosnakovski, D., Xu, Z., Li, W., Thet, S., Cleaver, O., Perlingeiro, R.C., and Kyba, M. (2008). Prospective Isolation of Skeletal Muscle Stem Cells with a Pax7 Reporter. Stem cells (Dayton, Ohio) 26, 3194–3024.

Brack, A.S., Bildsoe, H., and Hughes, S.M. (2005). Evidence that satellite cell decrement contributes to preferential decline in nuclear number from large fibres during murine age-related muscle atrophy. Journal of cell science 118, 4813–4821.

Brack, A.S., Conboy, M.J., Roy, S., Lee, M., Kuo, C.J., Keller, C., and Rando, T.A. (2007). Increased Wnt signaling during aging alters muscle stem cell fate and increases fibrosis. Science 317, 807–810.

Carlson, M.E., and Conboy, I.M. (2007). Loss of stem cell regenerative capacity within aged niches. Aging Cell 6, 371–382.

Chakkalakal, J.V., Jones, K.M., Basson, M.A., and Brack, A.S. (2012). The aged niche disrupts muscle stem cell quiescence. Nature 490.

Collins, B.C., Mader, T.L., Cabelka, C.A., Iñigo, M.R., Spangenburg, E.E., and Lowe, D.A. (2018). Deletion of estrogen receptor alpha in skeletal muscle results in impaired contractility in female mice. Journal of Applied Physiology 0, ull.

Conboy, I.M., Conboy, M.J., Wagers, A.J., Girma, E.R., Weissman, I.L., and Rando, T.A. (2005). Rejuvenation of aged progenitor cells by exposure to a young systemic environment. Nature 433, 760–764.

Conboy, I.M., and Rando, T.A. (2002). The regulation of Notch signaling controls satellite cell activation and cell fate determination in postnatal myogenesis. Dev Cell 3, 397–409.

Cosgrove, B.D., Gilbert, P.M., Porpiglia, E., Mourkioti, F., Lee, S.P., Corbel, S.Y., Llewellyn, M.E., Delp, S.L., and Blau, H.M. (2014). Rejuvenation of the muscle stem cell population restores strength to injured aged muscles. Nat Med 20, 255–264.

Dall, G.V., Hawthorne, S., Seyed-Razavi, Y., Vieusseux, J., Wu, W., Gustafsson, J.A., Byrne, D., Murphy, L., Risbridger, G., and Britt, K. (2018). Estrogen receptor subtypes dictate the proliferative nature of the mammary gland. The Journal of endocrinology.

Deschamps, A.M., Murphy, E., and Sun, J. (2010). Estrogen receptor activation and cardioprotection in ischemia reperfusion injury. Trends in cardiovascular medicine 20, 73–78.

Dumont, N.A., Wang, Y.X., and Rudnicki, M.A. (2015). Intrinsic and extrinsic mechanisms regulating satellite cell function. Development 142, 1572–1581.

Enns, D.L., Iqbal, S., and Tiidus, P.M. (2008). Oestrogen receptors mediate oestrogen-induced increases in post-exercise rat skeletal muscle satellite cells. Acta Physiologica 194, 81–93.

Enns, D.L., and Tiidus, P.M. (2008). Estrogen influences satellite cell activation and proliferation following downhill running in rats. J Appl Physiol 104, 347–353.

Fry, C.S., Lee, J.D., Mula, J., Kirby, T.J., Jackson, J.R., Liu, F., Yang, L., Mendias, C.L., Dupont-Versteegden, E.E., McCarthy, J.J., et al. (2015). Inducible depletion of satellite cells in adult, sedentary mice impairs muscle regenerative capacity without affecting sarcopenia. Nat Med 21, 76–80.

Fry, C.S., Porter, C., Sidossis, L.S., Nieten, C., Reidy, P.T., Hundeshagen, G., Mlcak, R., Rasmussen, B.B., Lee, J.O., Suman, O.E., et al. (2016). Satellite cell activation and apoptosis in skeletal muscle from severely burned children. The Journal of physiology 594, 5223–5236.

Fukada, S., Uezumi, A., Ikemoto, M., Masuda, S., Segawa, M., Tanimura, N., Yamamoto, H., Miyagoe-Suzuki, Y., and Takeda, S. (2007). Molecular signature of quiescent satellite cells in adult skeletal muscle. Stem cells (Dayton, Ohio) 25, 2448–2459.

Gelsomino, L., Panza, S., Giordano, C., Barone, I., Gu, G., Spina, E., Catalano, S., Fuqua, S., and Ando, S. (2018). Mutations in the Estrogen Receptor Alpha Hormone Binding Domain Promote Stem Cell Phenotype through Notch Activation in Breast Cancer Cell Lines. Cancer letters.

Greising, S.M., Baltgalvis, K.A., Kosir, A.M., Moran, A.L., Warren, G.L., and Lowe, D.A. (2011). Estradiol’s beneficial effect on murine muscle function is independent of muscle activity. J Appl Physiol 110, 109–115.

Greising, S.M., Baltgalvis, K.A., Lowe, D.A., and Warren, G.L. (2009). Hormone therapy and skeletal muscle strength: a meta-analysis. J Gerontol A Biol Sci Med Sci 64, 1071–1081.

Guo, L., Xu, J., Qi, J., Zhang, L., Wang, J., Liang, J., Qian, N., Zhou, H., Wei, L., and Deng, L. (2013). MicroRNA-17-92a upregulation by estrogen leads to Bim targeting and inhibition of osteoblast apoptosis. Journal of cell science 126, 978–988.

Hamilton, D.J., Minze, L.J., Kumar, T., Cao, T.N., Lyon, C.J., Geiger, P.C., Hsueh, W.A., and Gupte, A.A. (2016). Estrogen receptor alpha activation enhances mitochondrial function and systemic metabolism in high-fat-fed ovariectomized mice. Physiol Rep 4.

Harlow, S.D., Gass, M., Hall, J.E., Lobo, R., Maki, P., Rebar, R.W., Sherman, S., Sluss, P.M., and de Villiers, T.J. (2012). Executive summary of the Stages of Reproductive Aging Workshop + 10: addressing the unfinished agenda of staging reproductive aging. Menopause (New York, Ny) 19, 387–395.

Hindi, S.M., and Kumar, A. (2016). TRAF6 regulates satellite stem cell self-renewal and function during regenerative myogenesis. J Clin Invest 126, 151–168.

Hirai, H., Verma, M., Watanabe, S., Tastad, C., Asakura, Y., and Asakura, A. (2010). MyoD regulates apoptosis of myoblasts through microRNA-mediated down-regulation of Pax3. J Cell Biol 191, 347–365.

Kamanga-Sollo, E., White, M.E., Hathaway, M.R., Weber, W.J., and Dayton, W.R. (2010). Effect of Estradiol-17beta on protein synthesis and degradation rates in fused bovine satellite cell cultures. Domestic animal endocrinology 39, 54–62.

Keefe, A.C., Lawson, J.A., Flygare, S.D., Fox, Z.D., Colasanto, M.P., Mathew, S.J., Yandell, M., and Kardon, G. (2015). Muscle stem cells contribute to myofibres in sedentary adult mice. Nature communications 6, 7087.

Kim, J.H., Han, G.C., Seo, J.Y., Park, I., Park, W., Jeong, H.W., Lee, S.H., Bae, S.H., Seong, J., Yum, M.K., et al. (2016). Sex hormones establish a reserve pool of adult muscle stem cells. Nature cell biology 18, 930–940.

Kim, S., Ko, Y., Lee, H.J., and Lim, J.E. (2018). Menopausal hormone therapy and the risk of breast cancer by histological type and race: a meta-analysis of randomized controlled trials and cohort studies. Breast cancer research and treatment.

Kosir, A.M., Mader, T.L., Greising, A.G., Novotny, S.A., Baltgalvis, K.A., and Lowe, D.A. (2015). Influence of ovarian hormones on strength loss in healthy and dystrophic female mice. Medicine and science in sports and exercise 47, 1177–1187.

Kovanen V, A.P., Kokko K, Finni T, Tarkka IM, Tammelin T, Kujala UM, Sipilä S, Laakkonen EK (Submitted). Menopausal differences in blood count are associated with estrogen and follicle stimulating hormone: study protocol and novel result of the Estrogenic Regulation of Muscle Apoptosis (ERMA) cohort study. Biology of sex differences.

Kuang, S., Kuroda, K., Le Grand, F., and Rudnicki, M.A. (2007). Asymmetric Self-Renewal and Commitment of Satellite Stem Cells in Muscle. Cell 129, 999–1010.

La Colla, A., Pronsato, L., Milanesi, L., and Vasconsuelo, A. (2015). 17beta-Estradiol and testosterone in sarcopenia: Role of satellite cells. Ageing Res Rev 24, 166–177.

La Colla, A., Vasconsuelo, A., Milanesi, L., and Pronsato, L. (2016). 17beta-Estradiol Protects Skeletal Myoblasts from Apoptosis through P53, BCL-2 and FoxO Families. J Cell Biochem.

Laakkonen, E.K., Kulmala, J., Aukee, P., Hakonen, H., Kujala, U.M., Lowe, D.A., Kovanen, V., Tammelin, T., and Sipilä, S. (2017). Female reproductive factors are associated with objectively measured physical activity in middle-aged women. PLOS ONE 12, e0172054.

Lai, S., Collins, B.C., Colson, B.A., Kararigas, G., and Lowe, D.A. (2016). Estradiol modulates myosin regulatory light chain phosphorylation and contractility in skeletal muscle of female mice. Am J Physiol Endocrinol Metab 310, E724–733.

Le, G., Jergenson, M., Lowe, D., and Warren, G. (2015). Estradiol enhances neutrophil infiltration into traumatically-injured skeletal muscle. The FASEB Journal 29.

Le, G., Warren, G.L., and Lowe, D.A. (2017). 17b-Estradiol Rescues Low Force Potentiation in Ovariectomized Mice In Vivo. The FASEB Journal 31, 880.885.

Lowe, D.A., Warren, G.L., Ingalls, C.P., Boorstein, D.B., and Armstrong, R.B. (1995). Muscle function and protein metabolism after initiation of eccentric contraction-induced injury. J Appl Physiol 79, 1260–1270.

MacNeil, L.G., Baker, S.K., Stevic, I., and Tarnopolsky, M.A. (2011). 17beta-estradiol attenuates exercise-induced neutrophil infiltration in men. American journal of physiology Regulatory, integrative and comparative physiology 300, R1443–1451.

McClung, J.M., Davis, J.M., and Carson, J.A. (2007). Ovarian hormone status and skeletal muscle inflammation during recovery from disuse in rats. Exp Physiol 92, 219–232.

McClung, J.M., Davis, J.M., Wilson, M.A., Goldsmith, E.C., and Carson, J.A. (2006). Estrogen status and skeletal muscle recovery from disuse atrophy. J Appl Physiol 100, 2012–2023.

Moran, A.L., Nelson, S.A., Landisch, R.M., Warren, G.L., and Lowe, D.A. (2007). Estradiol replacement reverses ovariectomy-induced muscle contractile and myosin dysfunction in mature female mice. J Appl Physiol 102, 1387–1393.

Murphy, M.M., Keefe, A.C., Lawson, J.A., Flygare, S.D., Yandell, M., and Kardon, G. (2014). Transiently active Wnt/beta-catenin signaling is not required but must be silenced for stem cell function during muscle regeneration. Stem Cell Reports 3, 475–488.

Murphy, M.M., Lawson, J.A., Mathew, S.J., Hutcheson, D.A., and Kardon, G. (2011). Satellite cells, connective tissue fibroblasts and their interactions are crucial for muscle regeneration. Development 138, 3625–3637.

Nelson, J.F., Felicio, L.S., Randall, P.K., Sims, C., and Finch, C.E. (1982). A longitudinal study of estrous cyclicity in aging C57BL/6J mice: I. Cycle frequency, length and vaginal cytology. Biol Reprod 27, 327–339.

Nelson, L.R., and Bulun, S.E. (2001). Estrogen production and action. Journal of the American Academy of Dermatology 45, S116–S124.

Pallafacchina, G., Blaauw, B., and Schiaffino, S. (2013). Role of satellite cells in muscle growth and maintenance of muscle mass. Nutr Metab Cardiovasc Dis 23 Suppl 1, S12–18.

Phillips, S.K., Rook, K.M., Siddle, N.C., Bruce, S.A., and Woledge, R.C. (1993). Muscle weakness in women occurs at an earlier age than in men, but strength is preserved by hormone replacement therapy. Clin Sci (Lond) 84, 95–98.

Phillips, S.K., Sanderson, A.G., Birch, K., Bruce, S.A., and Woledge, R.C. (1996). Changes in maximal voluntary force of human adductor pollicis muscle during the menstrual cycle. The Journal of physiology 496 (Pt 2), 551–557.

Qaisar, R., Renaud, G., Hedstrom, Y., Pollanen, E., Ronkainen, P., Kaprio, J., Alen, M., Sipila, S., Artemenko, K., Bergquist, J., et al. (2013). Hormone replacement therapy improves contractile function and myonuclear organization of single muscle fibres from postmenopausal monozygotic female twin pairs. The Journal of physiology 591, 2333–2344.

Rader, E.P., and Faulkner, J.A. (2006). Effect of aging on the recovery following contraction-induced injury in muscles of female mice. J Appl Physiol 101, 887–892.

Ribas, V., Drew, B.G., Zhou, Z., Phun, J., Kalajian, N.Y., Soleymani, T., Daraei, P., Widjaja, K., Wanagat, J., de Aguiar Vallim, T.Q., et al. (2016). Skeletal muscle action of estrogen receptor alpha is critical for the maintenance of mitochondrial function and metabolic homeostasis in females. Sci Transl Med 8, 334ra354.

Ribas, V., Nguyen, M.T., Henstridge, D.C., Nguyen, A.K., Beaven, S.W., Watt, M.J., and Hevener, A.L. (2010). Impaired oxidative metabolism and inflammation are associated with insulin resistance in ERalpha-deficient mice. Am J Physiol Endocrinol Metab 298, E304–319.

Ronda, A.C., Vasconsuelo, A., and Boland, R. (2013). 17beta-estradiol protects mitochondrial functions through extracellular-signal-regulated kinase in C2C12 muscle cells. Cell Physiol Biochem 32, 1011–1023.

Sajko, Š., Kubínová, L., Cvetko, E., Kreft, M., Wernig, A., and Eržen, I. (2004). Frequency of M-Cadherin-stained Satellite Cells Declines in Human Muscles During Aging. Journal of Histochemistry & Cytochemistry 52, 179–185.

Sambasivan, R., Yao, R., Kissenpfennig, A., Van Wittenberghe, L., Paldi, A., Gayraud-Morel, B., Guenou, H., Malissen, B., Tajbakhsh, S., and Galy, A. (2011). Pax7-expressing satellite cells are indispensable for adult skeletal muscle regeneration. Development 138, 3647–3656.

Sanchez, A.M., Candau, R.B., and Bernardi, H. (2014). FoxO transcription factors: their roles in the maintenance of skeletal muscle homeostasis. Cell Mol Life Sci 71, 1657–1671.

Schultz, E., Jaryszak, D.L., Gibson, M.C., and Albright, D.J. (1986). Absence of exogenous satellite cell contribution to regeneration of frozen skeletal muscle. Journal of muscle research and cell motility 7, 361–367.

Severson, T.M., Kim, Y., Joosten, S.E.P., Schuurman, K., van der Groep, P., Moelans, C.B., Ter Hoeve, N.D., Manson, Q.F., Martens, J.W., van Deurzen, C.H.M., et al. (2018). Characterizing steroid hormone receptor chromatin binding landscapes in male and female breast cancer. Nature communications 9, 482.

Shea, K.L., Xiang, W., LaPorta, V.S., Licht, J.D., Keller, C., Basson, M.A., and Brack, A.S. (2010). Sprouty1 regulates reversible quiescence of a self-renewing adult muscle stem cell pool during regeneration. Cell Stem Cell 6, 117–129.

Shefer, G., Van de Mark, D.P., Richardson, J.B., and Yablonka-Reuveni, Z. (2006). Satellite-cell pool size does matter: defining the myogenic potency of aging skeletal muscle. Dev Biol 294, 50–66.

Sousa-Victor, P., Gutarra, S., Garcia-Prat, L., Rodriguez-Ubreva, J., Ortet, L., Ruiz-Bonilla, V., Jardi, M., Ballestar, E., Gonzalez, S., Serrano, A.L., et al. (2014). Geriatric muscle stem cells switch reversible quiescence into senescence. Nature 506.

Taaffe, D.R., Sipilä, S., Cheng, S., Puolakka, J., Toivanen, J., and Suominen, H. (2005). The effect of hormone replacement therapy and/or exercise on skeletal muscle attenuation in postmenopausal women: a yearlong intervention. Clinical Physiology and Functional Imaging 25, 297–304.

Tiidus, P.M., Holden, D., Bombardier, E., Zajchowski, S., Enns, D., and Belcastro, A. (2001). Estrogen effect on post-exercise skeletal muscle neutrophil infiltration and calpain activity. Can J Physiol Pharmacol 79, 400–406.

Troy, A., Cadwallader, Adam B., Fedorov, Y., Tyner, K., Tanaka, Kathleen K., and Olwin, Bradley B. (2012). Coordination of Satellite Cell Activation and Self-Renewal by Par-Complex-Dependent Asymmetric Activation of p38α/β MAPK. Cell Stem Cell 11, 541–553.

Vasconsuelo, A., Milanesi, L., and Boland, R. (2008). 17Beta-estradiol abrogates apoptosis in murine skeletal muscle cells through estrogen receptors: role of the phosphatidylinositol 3-kinase/Akt pathway. The Journal of endocrinology 196, 385–397.

Vasconsuelo, A., Milanesi, L., and Boland, R. (2010). Participation of HSP27 in the antiapoptotic action of 17beta-estradiol in skeletal muscle cells. Cell Stress Chaperones 15, 183–192.

Verdijk, L.B., Dirks, M.L., Snijders, T., Prompers, J.J., Beelen, M., Jonkers, R.A., Thijssen, D.H., Hopman, M.T., and Van Loon, L.J. (2012). Reduced satellite cell numbers with spinal cord injury and aging in humans. Medicine and science in sports and exercise 44, 2322–2330.

Verdijk, L.B., Snijders, T., Drost, M., Delhaas, T., Kadi, F., and van Loon, L.J. (2014). Satellite cells in human skeletal muscle; from birth to old age. Age (Dordrecht, Netherlands) 36, 545–547.

Wood, G.A., Fata, J.E., Watson, K.L.M., and Khokha, R. (2007). Circulating hormones and estrous stage predict cellular and stromal remodeling in murine uterus. Reproduction 133, 1035–1044.

